# Glycosaminoglycan content of a mineralized collagen scaffold promotes mesenchymal stem cell secretion of factors to modulate angiogenesis and monocyte differentiation

**DOI:** 10.1101/2021.03.23.436487

**Authors:** Marley J. Dewey, Vasiliki Kolliopoulos, Mai T. Ngo, Brendan A.C. Harley

## Abstract

Effective design of biomaterials to aid regenerative repair of craniomaxillofacial (CMF) bone defects requires approaches that modulate the complex interplay between exogenously added progenitor cells and cells in the wound microenvironment, such as osteoblasts, osteoclasts, endothelial cells, and immune cells. We are exploring the role of the glycosaminoglycan (GAG) content in a class of mineralized collagen scaffolds recently shown to promote osteogenesis and healing of craniofacial bone defects. We previously showed that incorporating chondroitin-6-sulfate or heparin improved mineral deposition by seeded human mesenchymal stem cells (hMSCs). Here, we examine the effect of varying scaffold GAG content on hMSC behavior, and their ability to modulate osteoclastogenesis, vasculogenesis, and the immune response. We report the role of hMSC-conditioned media produced in scaffolds containing chondroitin-6-sulfate (CS6), chondroitin-4-sulfate (CS4), or heparin (Heparin) GAGs on endothelial tube formation and monocyte differentiation. Notably, endogenous production by hMSCs within Heparin scaffolds most significantly inhibits osteoclastogenesis via secreted osteoprotegerin (OPG), while the secretome generated by CS6 scaffolds reduced pro-inflammatory immune response and increased endothelial tube formation. All conditioned media down-regulated many pro- and anti-inflammatory cytokines, such as IL6, IL-1β, and CCL18 and CCL17 respectively. Together, these findings demonstrate that modifying mineralized collagen scaffold GAG content can both directly (hMSC activity) and indirectly (production of secreted factors) influence overall osteogenic potential and mineral biosynthesis as well as angiogenic potential and monocyte differentiation towards osteoclastic and macrophage lineages. Scaffold GAG content is therefore a powerful stimulus to modulate reciprocal signaling between multiple cell populations within the bone healing microenvironment.

## 1. Introduction

Over 2 million bone graft surgeries occur annually worldwide, including over 500,000 in the US alone, with an estimated cost burden of 3.9 billion USD [1-4]. Critical-sized craniomaxillofacial (CMF) defects commonly result from tumor excisions, high-energy impacts in both the civilian (e.g., sports injury, motor vehicle accident) and Warfighter (e.g., penetrating battlefield injuries), as well as congenital disorders [5-7]. Fracture healing and bone remodeling is a complex mechanism that involves the interplay and crosstalk of multiple cell types including osteoblasts, osteoclasts, endothelial cells, and immune cells. Fracture healing can be broadly categorized in three phases: pro-inflammatory phase, proliferative phase, and remodeling phase [8]. The pro-inflammatory phase occurs immediately after the fracture with the formation of a hematoma and the recruitment of immune cells such as monocytes and macrophages. Macrophages, which are derived from monocytes, are phagocytic cells that regulate inflammation and early vascularization to further promote cell recruitment. Macrophages have been largely characterized as either pro-inflammatory (M1) or anti-inflammatory (M2). M1 macrophages take part in the pro-inflammatory phase and have been shown to regulate mesenchymal stem cells (MSCs) and promote osteogenesis [8-11]. M2 macrophages are associated with the proliferative phase by promoting collagen deposition and producing wound healing factors such as interleukin 10 (IL-10), bone morphogenetic protein 2 (BMP-2), and vascular endothelial growth factor (VEGF), thereby stimulating tissue homeostasis [12]. Disruptions to the immune response can disrupt bone regeneration by inducing a persistence of inflammatory stimuli, which can lead to fibrous tissue surrounding the implant and inhibiting healing [9-11].

Multiple signaling axes may contribute to improved bone regeneration beyond the inflammatory axes. In particular, vascularization plays a vital role during the proliferative phase, as the fracture site lacks in oxygen and nutrients, and new vessels can provide surrounding cells with waste removal, oxygen, and nutrients. VEGF is a key soluble factor involved in angiogenesis and enhances proliferation and development of endothelial cells as well as the differentiation of primary osteoblast cells [8]. Further, hMSC-secreted VEGF has been shown to stimulate angiogenesis through endothelial cell migration, proliferation, and differentiation [13]. Finally, during bone remodeling, monocytes can also produce osteoclasts that act in concert with osteoblasts to maintain an equilibrium bone mass. While osteoclast progenitors express receptor activator of nuclear factor kappa-Β (RANK), which binds to osteoblast-associated receptor activator of nuclear factor kappa-Β ligand (RANKL) and signals for osteoclast differentiation and bone resorption, hMSCs are able to produce a soluble decoy receptor called osteoprotegerin (OPG) that binds RANKL and inhibits RANK signaling to reduce osteoclast activity [14-17]. An increase in secreted OPG has been linked to less resorption by osteoclasts on mineralized collagen scaffolds and provides a route to improve early stage graft integration with the surrounding wound site and improved quantity and quality of bone repair [14, 15].

Due to their size and often irregular shape, CMF bone defects are generally reconstructed with autografts, allografts, or biomaterial alternatives [18-21]. Autografts are considered the gold standard as they lead to osteoconduction, osteoinduction and osteogenesis, but they are limited due to supply and donor site morbidity [22]. Allografts are the second most commonly used as they are widely available, but are associated with risks of disease transmission and quality of healing [23, 24]. Regenerative biomaterials seek to overcome these limitations, and a subclass of promising materials are those based on mineralized collagen scaffold technologies. These scaffolds consist of elements commonly found in bone such as type I collagen, calcium and phosphate ions, and glycosaminoglycans. Mineralized collagen scaffolds are highly porous and have been shown to induce mineral formation *in vitro* without the use of osteogenic supplements, as well as produce bone *in vivo* [25-34]. Efforts to improve the quality and speed of repair are beginning to examine the role of additional inclusions within the collagen scaffold, such as glycosaminoglycans (GAGs). GAGs have been shown to play an important role in bone homeostasis [35]. Collagen-GAG scaffolds have been described and characterized extensively in literature using various methods including tensile and compressive tests, cell adhesion and migration behaviors within the material [36-44]. Recently, inclusion of chondroitin-4-sulfate (CS4), chondroitin-6-sulfate (CS6), and heparin (Heparin) have been investigated for their role in bone formation in mineralized collagen scaffolds [45-49]. Although CS6 and Heparin demonstrated improved mineral biosynthesis, successful healing of CMF defects does not solely depend on mineral deposition but also on the ability to modulate the immune response and induce angiogenesis. Heparin and heparan sulfate have been shown to enhance angiogenesis and induce vascularization while mitigating inflammation [50-54]. Chondroitin sulfates (CS) display anti-inflammatory properties by suppressing the nuclear translocation of nuclear factor-κB (NF-κB); however, they are not known to induce angiogenesis [51-54]. Thus, our objective was to study the impact of scaffold glycosaminoglycan content on processes linked to immune response, bone resorption, and vascular formation.

In this work, we modify the GAG content in our mineralized collagen scaffolds to define the effect of including CS6, CS4, or Heparin within the scaffold microenvironment on downstream processes associated with angiogenesis, osteoclastogenesis, and the immune response. We examine the effects of GAG-dependent changes in hMSC conditioned media on angiogenesis through the use of a Matrigel tube formation assay, and on immune response by measuring monocyte differentiation towards osteoclast versus macrophage lineage over 21 days of *in vitro* cell culture. We believe that modulating scaffold GAG content provides a powerful indirect stimuli, mediated by MSC-driven production of factors on a broad population of cells involved in bone repair (monocytes, macrophages, osteoclasts, and endothelial cells).

## 2. Materials and Methods

### 2.1 Experimental design

The goal of this study was to determine the impact of glycosaminoglycans within mineralized collagen scaffolds on angiogenesis and the immune response (**Fig. 1**). Mineralized collagen scaffolds containing glycosaminoglycans CS6, CS4, or Heparin were cultured with human mesenchymal stem cells for 21 days. The media was collected and pooled after the first 6 days for a vascular tube formation assay, in which media was added to human umbilical vein endothelial cells (HUVECs) seeded on top of Matrigel and vessel network length quantified after 6 and 12hrs. Secretion of OPG and VEGF by hMSCs as a function of scaffold GAG content was quantified after 21 days of hMSC culture. Further, hMSC-GAG conditioned media was added to monocyte culture for a 21-day culture period to compare the effect of hMSCs-GAG conditioned media versus control media (monocyte RPMI media) on monocyte specification. Macrophage versus osteoclast specification by monocytes in response to conditioned media generated by hMSCs seeded within scaffolds fabricated using different GAGs was assessed via a combination of gene expression as well as functional assays including cytokine arrays, targeted ELISAs, and Tartrate-Resistant Acid Phosphatase (TRAP) staining.

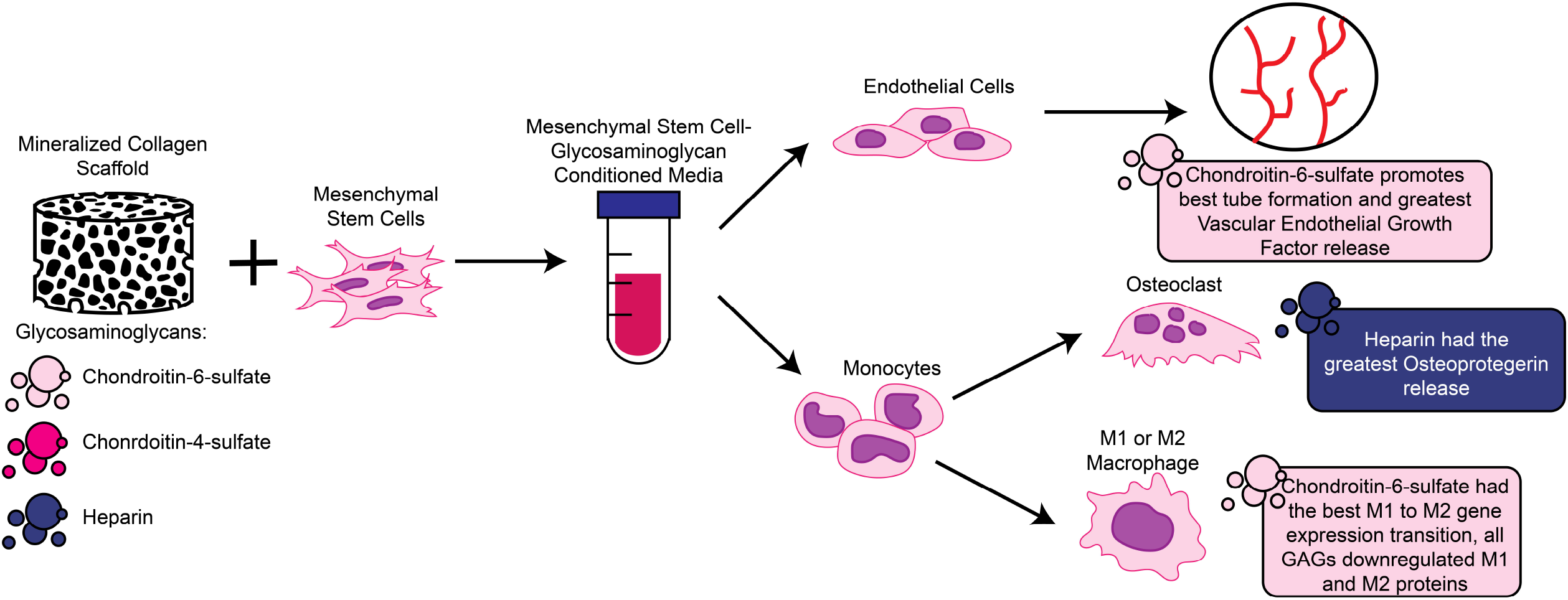

### 2.2 Mineralized collagen-glycosaminoglycan scaffold fabrication

Mineralized collagen-glycosaminoglycan scaffolds were fabricated via lyophilization from mineralized collagen precursor suspensions with varying glycosaminoglycans as previously described [26, 31, 55]. Briefly, type I bovine collagen (1.9 w/v% Sigma-Aldrich, Missouri USA), calcium salts (calcium hydroxide and calcium nitrate tetrahydrate, Sigma-Aldrich), and glycosaminoglycans (0.84 w/v%) were homogenized in mineral buffer solution (0.1456 M phosphoric acid/0.037 M calcium hydroxide). Glycosaminoglycans evaluated in this study included chondroitin-6-sulfate (CS6, Chondroitin sulfate sodium salt from shark cartilage, CAS Number 9082-07-9, Sigma-Aldrich), chondroitin-4-sulfate (CS4, Bovine-derived, Sodium Chondroitin Sulfate A, CAS Number 39455-18-0, Toronto Research Chemicals, Ontario, Canada), or heparin (Heparin, Heparin sodium salt from porcine intestinal mucosa, CAS 9041-08-1, Sigma-Aldrich). The collagen precursor suspensions were subsequently transferred to aluminum molds and lyophilized into scaffolds using a Genesis freeze-dryer (VirTis, Gardener, New York USA). The suspensions were cooled at a constant rate of 1 °C/min from 20 °C to - 10 °C followed by a hold at -10 °C for 2 hours. The frozen suspension was subsequently sublimated at 0 °C and 0.2 Torr, resulting in a porous scaffold network. After lyophilization, a 6 mm diameter biopsy punch (Integra LifeSciences, New Jersey, USA) was used to create individual scaffolds.

### 2.3 Sterilization, hydration, and scaffold crosslinking

All scaffolds were placed in sterilization pouches and sterilized via ethylene oxide treatment for 12 hrs using a AN74i Anprolene gas sterilizer (Andersen Sterilizers Inc., Haw River, North Carolina USA) [25]. After sterilization, all subsequent steps proceeded using aseptic techniques. Sterile scaffolds were then hydrated and crosslinked using previously described EDC-NHS chemistry [26]. Briefly, scaffolds were soaked in 100% ethanol, then washed multiple times in phosphate buffered saline (PBS), followed by EDC-NHS crosslinking. Scaffolds were further washed in PBS and finally soaked in basal growth media for 48 hours prior to cell seeding.

### 2.4 Cell culture and conditioned media

#### 2.4.1 Human mesenchymal stem cell culture and scaffold seeding

Passage 4 human mesenchymal stem cells (hMSCs) (StemBioSys Inc, hBM-MSC, Lot# BM 17-002) were expanded to passage 5 using culture media containing low glucose DMEM and glutamine, 10% mesenchymal stem cell fetal bovine serum (Gemini, California, USA), and 1% antibiotic-antimycotic (Gibco, Massachusetts, USA) in an incubator at 37LC and 5% CO2. Mycoplasma contamination was tested with a MycoAlert™ Mycoplasma Detection Kit (Lonza, Walkersville, MD) prior to seeding on scaffolds. All experiments used cells that tested negative for mycoplasma. hMSCs were seeded on hydrated scaffolds at a density of 100,000 cells per scaffold using a previously described method [36, 56]. Briefly, half of the cells were seeded on the top of the scaffold and allowed to attach for 30 min in an incubator, followed by seeding on the bottom of the scaffold for a 1.5 hr incubation prior to adding basal media.

#### 2.4.2 Mesenchymal stem cell conditioned media collection and preparation

Media was collected from each hMSC-seeded scaffold and replaced every third day for 21 days. The media collected from all timepoints was combined into a batch for the respective scaffold group. Three variations of hMSC-GAG conditioned media were created (CS6, CS4, and Heparin conditioned media). A control media was used as a standard, which represents the hMSC basal media without scaffold and cell conditioning. Each media group was diluted at a 1:1 ratio with RPMI media.

#### 2.4.3 Human umbilical vein endothelial cell (HUVEC) culture

HUVECs (Lonza) were cultured in Endothelial Growth Medium 2 (EGM2) (Lonza) supplemented with plasmocin prophylactic (Invivogen, San Diego, CA) to prevent mycoplasma contamination. Cells were maintained at 37 °C and 5% CO_2_ and were used before passage 5.

#### 2.4.4 THP-1 monocyte cell culture and plate seeding

THP-1 monocyte cells (ATCC® TIB-202™, Maryland, USA) derived from human peripheral blood were used to determine the effect of hMSC/glycosaminoglycan conditioned media on macrophage and osteoclast differentiation. THP-1 cells were expanded in RPMI medium (ATCC® 30-2001™) supplemented with 10% heat inactivated fetal bovine serum (HI-FBS, Gibco, ThermoFisher Scientific, Massachusetts, USA), 100 U/ml penicillin and 100 μg/ml streptomycin sulfate (ThermoFisher Scientific). THP-1 cells were not allowed to exceed 0.8 million cells/ml during culture. THP-1 cells were centrifuged at 130 rcf for 5 min and resuspended to achieve a concentration of 20 million cells/ml. Cells were seeded on 24-well plates using 50 μl of the cell suspension equating to 1 million cells/well. Each well was then supplemented with 1 ml of each 1:1 hMSC conditioned media (control, CS6, CS4, or Heparin) and RPMI media.

### 2.5 Matrigel tube formation assay with hMSC conditioned media

100 µL of growth factor reduced Matrigel (Corning, Massachusetts, USA) was dispensed per well in a 96-well plate and allowed to gel at 37 °C for one hour. 10,000 HUVECs were mixed with 150 µL of each hMSC-conditioned media sample (CS4, CS6, Heparin) and pipetted over the Matrigel layer in each well (n=6) [57]. Unconditioned growth media (basal hMSC media) was used as a negative control. Tube formation was visualized and brightfield images were recorded using a Leica DMI4000 B (Leica Microsystems, Illinois, USA) at 6 and 12 hours. Three regions were imaged per well. The “Angiogenesis Analyzer” toolset on ImageJ (NIH, Maryland, USA) was used to quantify and compare tube formation between groups in terms of total network length [58]. For each sample, the reported length is the average from the three regions of interest.

### 2.6 Cell number quantification and cytotoxicity of monocytes

DNA content of monocytes was quantified using the Quant-iT™ PicoGreen ® dsDNA Assay kit (Invitrogen) according to the manufacturer’s instructions (n=5). The sample readings were quantified using a standard curve prepared with known DNA concentrations following the same procedure. At each of the timepoints (day 3, 7, 14, 21), all media was aspirated from the 2D wells and 50 μl of fresh media was added into each well. The cells were lightly scraped off the surface of the plate using a cell scraper (Corning, New York, USA). A 10 μl sample was taken from each well and added into 100 μl of 1x PicoGreen solution in TE (supplied from kit). Following a 5 min incubation, fluorescence readings (excitation 480 nm and emission 520 nm) were measured with a F200 spectrophotometer (Tecan, Männedorf, Switzerland). Media controls taken from unseeded scaffolds for each GAG type (n = 3) were used to subtract the background from the samples. The intensities were then converted to DNA concentrations using the slope and intercept extracted from the standard curve.

Cytotoxicity of monocytes cultured in hMSC-GAG media was quantified with an LDH-Glo™ Cytotoxicity Assay (Promega, Wisconsin, USA). Lactate dehydrogenase (LDH) is released from cells upon membrane damage, and this amount released into surrounding media can be used to quantify cytotoxicity. At each of the timepoints (day 3, 7, 14, 21), 5 μL of media surrounding monocytes was added to 95 μL LDH storage buffer and stored at -20°C until analysis (n=5). Media from monocytes prior to adding to wells with hMSC-GAG media was used as a control. 10 μL of sample was added to 40 μL LDH storage buffer to react with enzyme and reductase solutions and was read in duplicate for luminescence using a M200 spectrophotometer (Tecan) with a 1 second integration time. Luminescence readings were normalized to the control group, with a value of 1 representing the amount of LDH released from monocytes before seeding in hMSC-GAG media and values greater than 1 indicating greater release of LDH and possible cytotoxicity.

### 2.7 Monocyte gene expression

RNA was isolated from monocytes exposed to control, CS6, CS4, and Heparin scaffold conditioned media groups (n=5) across 21 days (Days 0, 3, 7, 14, 21) using a RNAqueous™-Micro Total RNA Isolation Kit (ThermoFisher Scientific). RNA was then reverse transcribed to cDNA using a QuantiTect Reverse Transcription Kit (Qiagen, Germany) and an S100 thermal cycler (Bio-Rad, California, USA). Samples for real-time PCR were prepared using 30 ng of cDNA in 20 μL reactions with Taqman fast advanced master mix and Taqman gene expression assays (Applied Biosystems, California, USA). All Taqman assays were pre-validated by the manufacturer. 96-well PCR plates were read using a Quantstudio 3 System (ThermoFisher Scientific) and results were analyzed using the delta-delta CT method, with gene expression expressed as fold change normalized to the housekeeping gene, β-Actin, as suggested by Maess et. al. [59]. Details of all genes examined are provided in **Supp. Table 1**.

### 2.8 Soluble factor analysis

#### 2.8.1 ELISAs of hMSC conditioned media

OPG and VEGF ELISAs were used on pooled hMSCs-GAG conditioned media after 3, 9, 15, and 21 days of cell culture to determine the amount of protein secreted by hMSCs cultured on mineralized collagen scaffolds with different glycosaminoglycans (n=6). ELISA kits were purchased from R&D Systems (Minnesota, USA). 25 μL of sample was used with 75 μL of reagent diluent during the sample binding step. Wells were read for absorbance with a spectrophotometer (M200, Tecan, Switzerland) and a non-conditioned normal growth media control was subtracted from the sample results.

#### 2.8.2 Cytokine array of monocyte conditioned media

A custom cytokine array from RayBiotech (Georgia, USA) was used to examine proteins cardiotrophin 1 (CT-1), platelet-derived growth factor homodimer (PDGF-BB), interferon gamma (IFN-gamma), tumor necrosis factor alpha (TNFa), interleukin 1 beta (IL-1beta), Vascular Endothelial Growth Factor A (VEGFA), interleukin 6 (IL-6), IL-10, C-C Motif Chemokine Ligand 18 (CCL18), C-C Motif Chemokine Ligand 22 (CCL22), Matrix metallopeptidase 9 (MMP9), C-C Motif Chemokine Ligand 17 (CCL17), interleukin 4 (IL-4), interleukin 13 (IL-13) in media surrounding monocytes pooled throughout 21 days of culture (n=3). A mixture of hMSC and RPMI media on monocytes was used as a control. Media was pooled from equal volumes at each timepoint to create a total volume of 1mL for each membrane (days 3, 6, 9, 12, 15, 18, 21). Cytokines were detected using a LAS 4010 luminescent image analyzer (GE Healthcare, New Jersey, USA), and membranes were normalized to each other using positive spots and analyzed with ImageJ. Samples of hMSC conditioned media were subtracted from monocyte conditioned media to quantify the cytokines released only by the monocytes. After subtraction, both hMSC conditioned media and monocyte conditioned media were normalized to the control media containing RPMI and hMSC media without conditioning to obtain a fold change.

### 2.9 Tartrate-Resistant Acid Phosphatase (TRAP) staining of monocytes

TRAP staining was performed on monocytes (n=6) after 14 and 21 days of culture in hMSC-GAG conditioned media following a protocol from the University of Rochester Medical Center (Center for Musculoskeletal Research) [60, 61]. Reagents were purchased from Sigma-Aldrich (Missouri, USA). Briefly, media was aspirated from wells and cells were washed once in distilled water before fixation for 5 minutes in Shandon™ Formal-Fixx™ 10% Neutral Buffered Formalin (ThermoFisher Scientific, Massachusetts, USA). Cells were washed again in distilled water, incubated in TRAP medium for 30 minutes at 37°C, and rinsed again in distilled water. 0.02% fast green was added for 30 seconds, and finally cells were rinsed again in distilled water and air dried. Once dry, each well was imaged with a Leica DFC295 (Leica, Wetzlar, Germany) microscope to identify osteoclasts (red) and other adherent cells (green/blue) (n=6). TRAP images were compared qualitatively to images of monocytes before washing and staining.

### 2.10 Statistics

Statistical analysis followed practices and procedures by Ott and Longnecker [62]. Statistical analysis utilized a 95% confidence interval for all tests. Data was first assessed for any outliers with a Grubbs test. After removal of outliers, normality of residuals was assessed with a Shapiro-Wilk test. Equal variance was assessed on residuals using the Levene’s Test. All groups underwent statistical testing to determine significance between groups depending on normality and equal variance assumptions. For normal data with equal variance, an ANOVA with a Tukey-post hoc was used. For non-normal data with equal variance, a Kruskal-Wallis test was used followed by a Dunn test using the Benjamini-Hochberg method. For normal data with unequal variance, a Welch’s ANOVA with a Games-Howell post-hoc was used. For non-normal data with unequal variance, a Welch’s Heteroscedastic F test with trimmed means and winsorized variances was used with a Games-Howell post-hoc. Sample numbers were based on previous analyses of similar experiments, and mentioned in figure captions [63-66]. Data is expressed as average ± standard deviation unless otherwise noted.

## 3. Results

### 3.1 MSC secretome is sensitive to the glycosaminoglycan content of the mineralized collagen GAG scaffold

Osteoprotegerin and VEGF ELISAs were used to determine the amount of protein released from hMSCs seeded on mineralized collagen scaffolds with various GAGs over the course of 21 days of culture. All groups demonstrated significant OPG production over 21 days, with no significant differences between groups before day 9. However, between 15-21 days of culture, hMSCs in mineralized collagen scaffolds containing Heparin produced significantly (p < 0.05) more OPG than in CS6 or CS4 scaffolds (**Fig. 2A**). Similarly, increased secretion of VEGF was observed by hMSCs in all scaffolds over 21 days, with significantly (p < 0.05) greater VEGF produced by the hMSCs in the CS6 group versus Heparin scaffolds between days 9 and 15 (**Fig. 2B**).

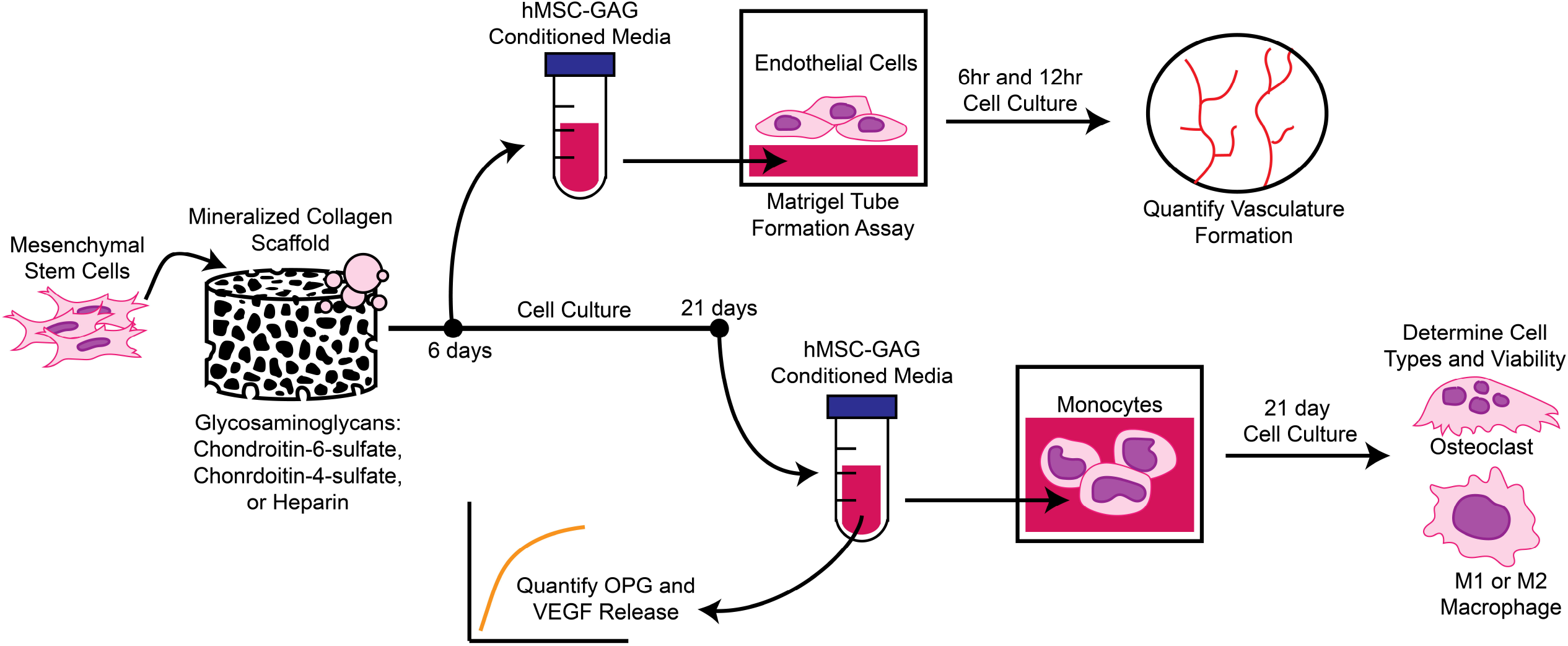

### 3.2 Secretome generated by MSCs in chondroitin-6-sulfate scaffolds promote greater vessel network formation than other glycosaminoglycans

A Matrigel tube formation assay was used to assess the angiogenic potential of proteins secreted by hMSCs over the first 6 days of culture. As rapidly as six hours post-seeding, a significant increase in tube formation was observed for all hMSC conditioned media compared to basal media (negative control), indicating a functional effect of angiogenic factors in all hMSC conditioned media (**Fig. 3**). Notably, quantification of endothelial cell network formation demonstrated that the magnitude of angiogenic potential was significantly influenced by scaffold GAG content, with a significant increase in total network length in response to hMSC secretome produced within CS6 containing scaffolds. This effect was maintained through twelve hours, with CS6 conditioned media promoting significantly increased total network length compared to the basal media control as well as CS4 and Heparin containing scaffolds (**Supp. Fig. 1**).

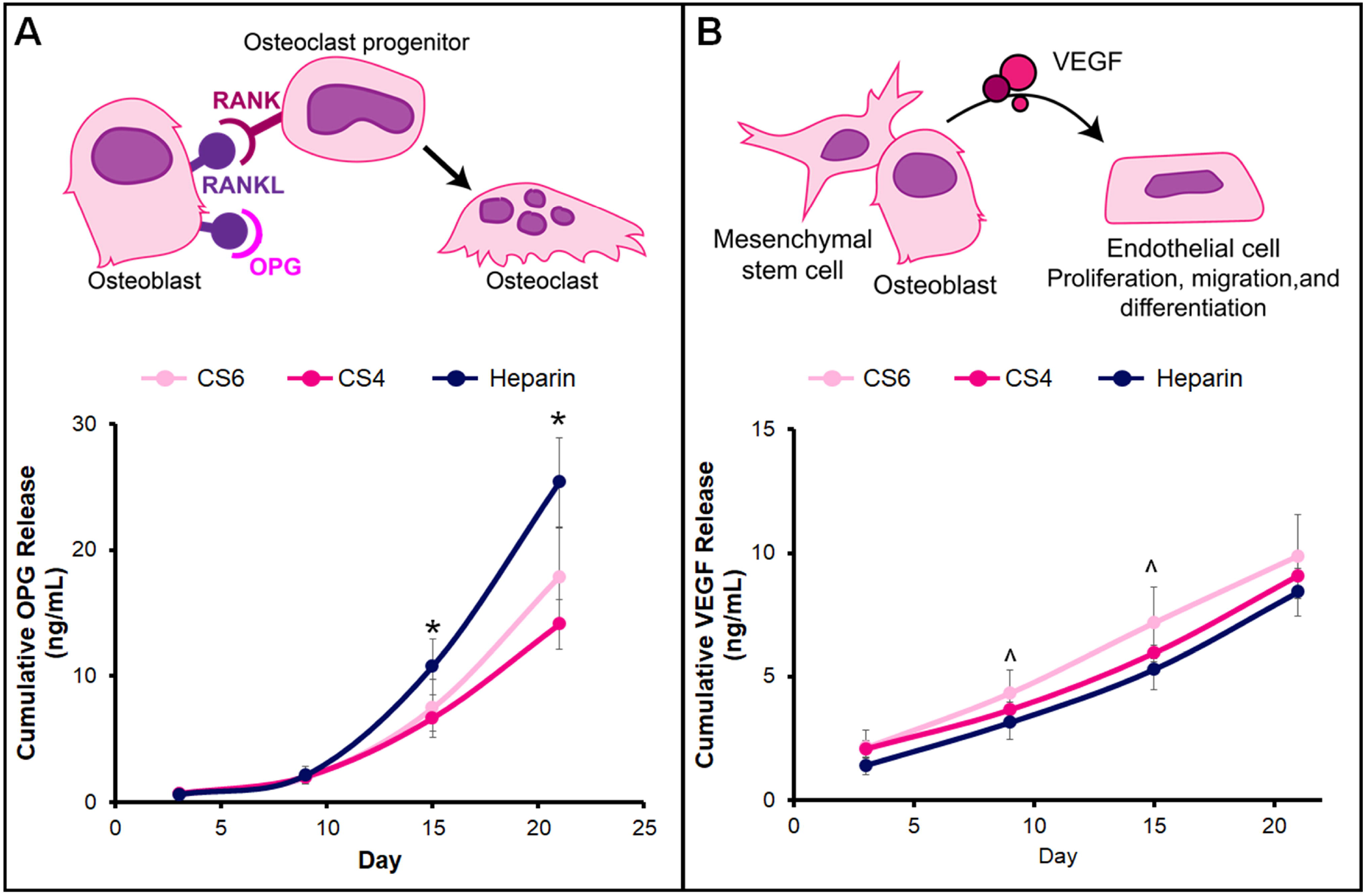

### 3.3 hMSC conditioned media impacts early monocyte cell health, but after 21 days there is no appreciable cytotoxic effect

We subsequently examined monocyte viability in hMSC conditioned media over 21 days. At day 7 and 14, all groups displayed a decrease in DNA content compared to day 0, and monocytes exposed to hMSC conditioned media in Heparin scaffolds had significantly more DNA content than all other groups at day 7. At day 21, all groups displayed an increase in DNA content; however, CS4 displayed significantly less DNA than the control (**Supp. Fig. 2A**). We then quantified monocyte cytotoxicity in response to hMSC conditioned media as a function of scaffold GAG content. All media groups including the control (hMSC and monocyte media), displayed LDH values above 1 by day 3, suggesting cell membrane damage and possible cell death (**Supp. Fig. 2B**). However, for the remainder of the culture LDH release from cells was reduced, suggesting less death or damage. At day 7, monocytes exposed to basal media (control) had significantly (p < 0.05) higher LDH release than any hMSC conditioned media, and results between groups were largely equivalent over the 21 day culture. This suggests factors produced endogenously by hMSCs in mineralized collagen scaffolds, regardless of GAG content, did not contribute to cell death, but rather basal hMSC media may have induced some cytotoxic effect on the monocytes.

### 3.4 Scaffold glycosaminoglycan content induces MSC-mediated immunosuppressive phenotype, with MSC secretome generated in CS6 scaffolds maximally inducing M1 to M2 gene expression transition

Gene and protein expression patterns were used to assess patterns of monocyte specification in response to MSC conditioned media for up to 21 days in culture. Broadly, a more M1-like phenotype was ascribed to increased expression of *IL-1*β and *TNF*α genes, with increased *CCL22* and *CCL17* used as M2a proxies and *CD163* and *MARCO* as M2c genes [67-70]. Although *IL-1*β was downregulated throughout the 21 days of the experiment, *TNF*α was upregulated starting at day 7 for the chondroitin groups whereas at day 21 we observe an increase in expression for CS4 and Heparin.(**Fig. 4**). Starting at day 14, *CCL22* was upregulated in CS6 compared to the media control. Although there were no significant differences with *CCL17* between groups, an increasing trend with time was observed. *CD163* was significantly (p < 0.05) downregulated by MSC secretome generated in CS6 and Heparin scaffolds at days 3, 7, and 21 while *MARCO* was significantly (p < 0.05) downregulated in response to MSC secretome generated in all groups on day 7 compared to the control.

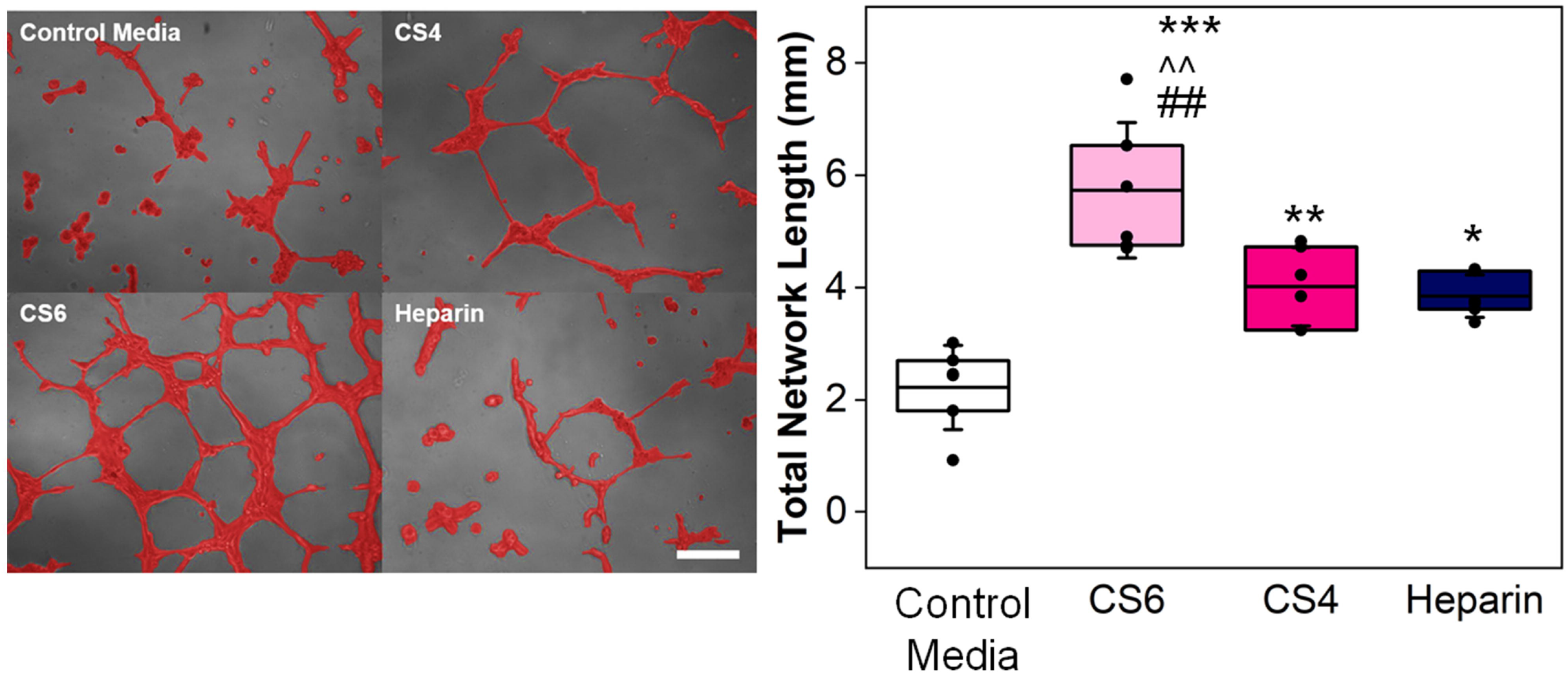

Patterns of monocyte differentiation were also characterized via cytokine array. IFNγ, TNFα, IL1β and IL6 cytokines are commonly ascribed to a more M1 phenotype [67, 70-73], while cytokine factors including IL4, IL13, CCL17, CCL18, CCL22, and MMP9 associated with a more M2 phenotype [67, 71-73]. We observed an overall downregulation of both M1 and M2 associated proteins in response to MSC secretome in CS6 scaffolds (**Fig. 5, Supp. Table 2 & 3**). Specifically, IFNγ, TNFα, and IL1β were all significantly (p < 0.05) downregulated. Furthermore, CCL18, CCL22, and IL4 were all significantly (p < 0.05) downregulated in all media groups while IL13 was significantly (p < 0.05) downregulated in the CS6 scaffold conditioned media. There was no significance between groups for IL10, MMP9 or CCL17, though all trended towards decreased expression as well. These results indicate an overall immunosuppressive effect of MSC conditioned media, with the most significant effects observed with the secretome generated by MSCs cultured in CS6-containing collagen scaffolds.

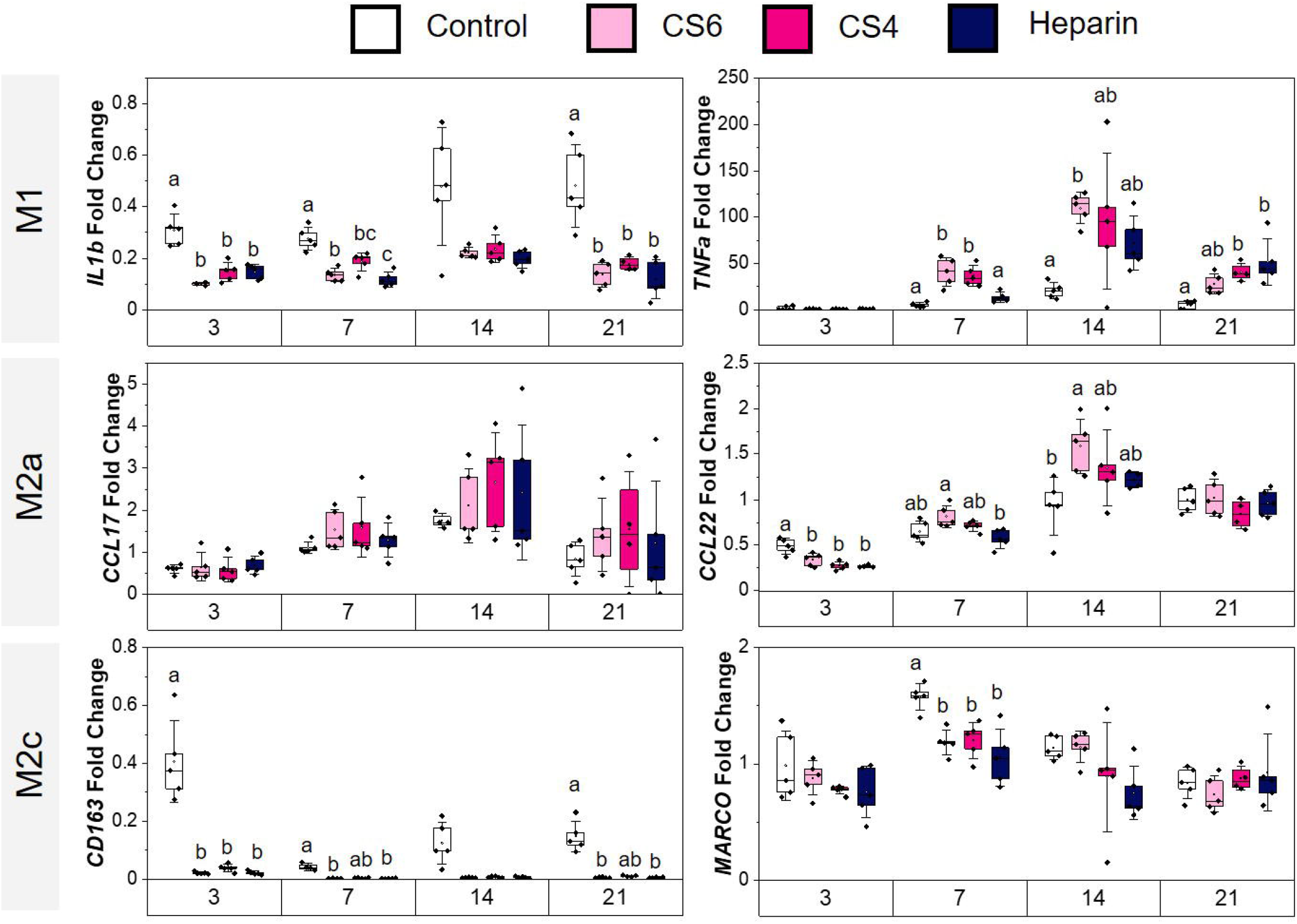

### 3.5 While CS6 scaffold conditioned media promotes late-stage osteoclast gene expression, MSC secretome generally downregulates osteoclast-associated protein expression

Gene (*SEMA4D, CTHRC1*) and protein (*CT-1, PDGF-BB)* expression patterns were subsequently used to assess osteoclast-related specification patterns in response to MSC conditioned media for up to 21 days in culture. Expression of *SEMA4D*, expressed exclusively by osteoclasts and known to inhibit bone formation [74], was significantly (p < 0.05) downregulated after 21 days in response to MSC secretome generated in all scaffolds groups (**Fig. 6A**). *CTHRC1* expression, upregulated in active osteoclasts and bone remodeling [74], was significantly downregulated in CS6 and Heparin groups at day 3 compared to the control (**Fig. 6A**). CT-1 and PGDF-BB are both expressed by osteoclasts, acting as a coupling factor with osteoblasts and osteocytes during the remodeling phase of healing and as a pre-regenerative factor (angiogenesis, osteogenic differentiation), respectively [74]. Generally, both CT-1 and PDGF-BB were expressed at a significantly (p < 0.05) reduced level in response to MSC secretome (**Fig. 6B**). There were no differences in PDGF-BB secretion by hMSCs on various GAG scaffolds; however, when added to monocytes, PDGF-BB was significantly (p < 0.05) upregulated in the monocytes without GAG conditioning (**Fig. 6, Supp. Table 2 & 3**).

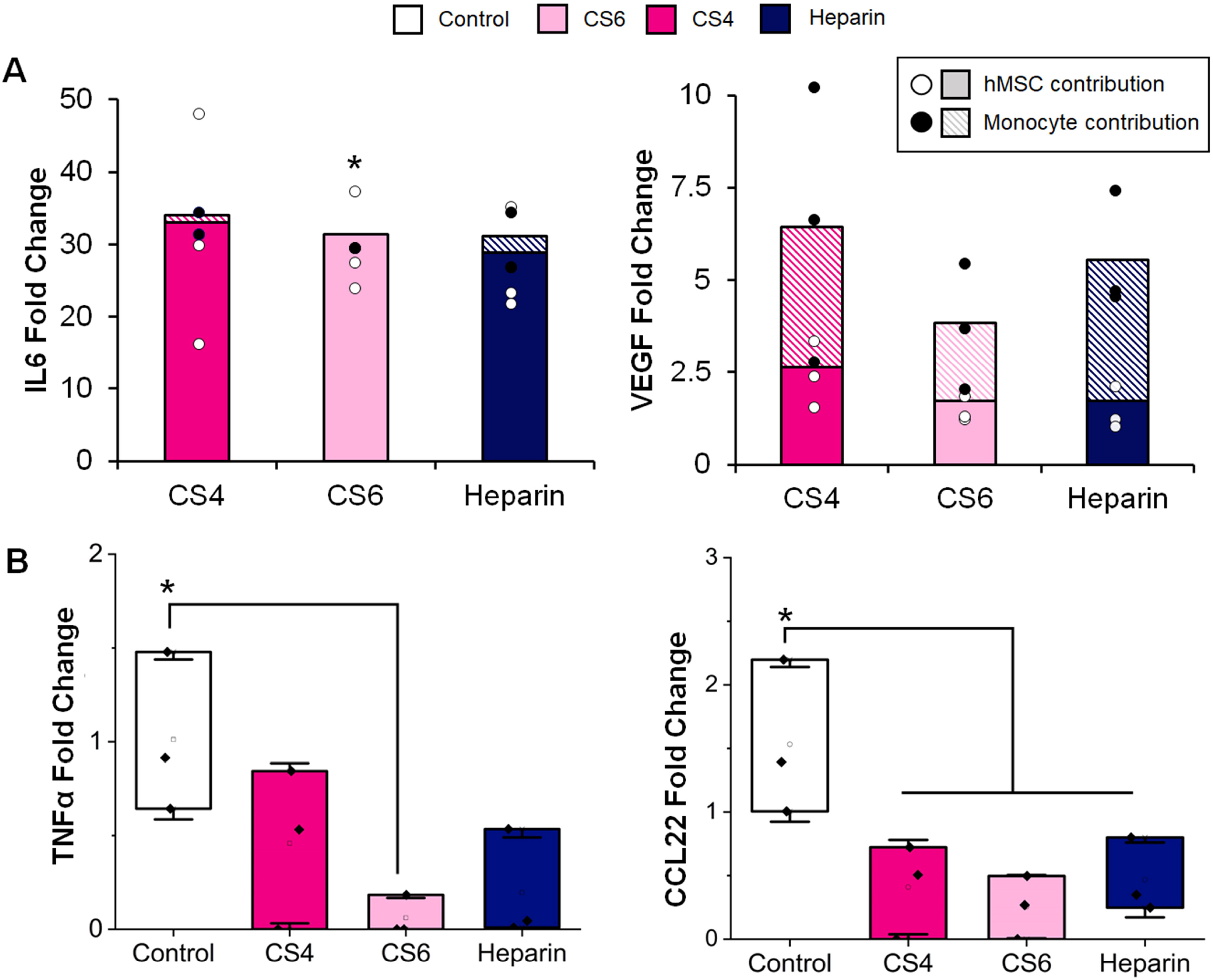

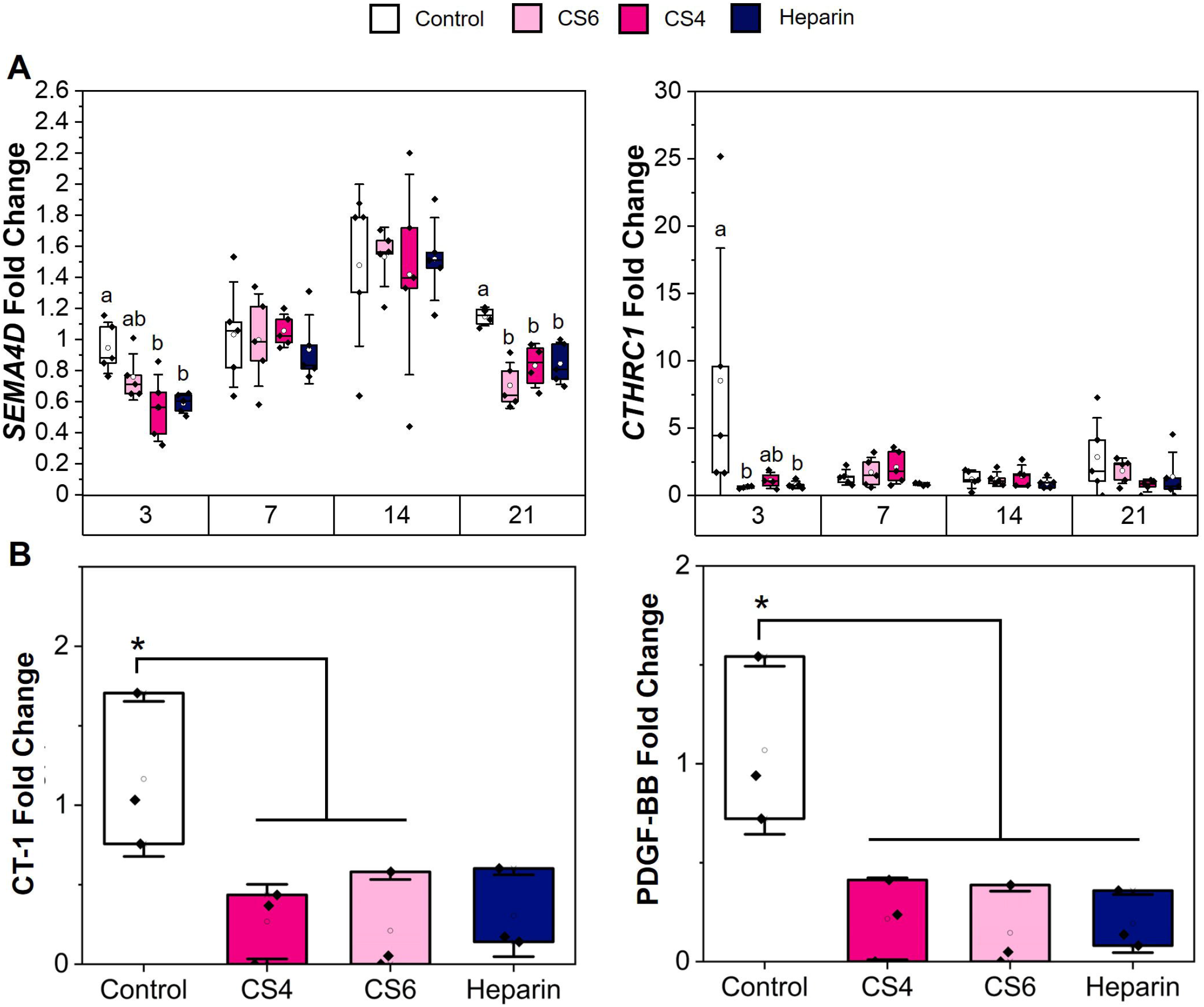

### 3.6 Adherent cell types were found in all groups, especially Heparin

TRAP staining was performed on monocytes cultured in hMSC-GAG media after 14 and 21 days of culture to stain for osteoclasts. No noticeable osteoclasts were found in any of the groups. By day 14 and day 21 there were very few attached cells in response to control media, but by day 14 there were noticeably more adherent cells in response to hMSC conditioned media for all scaffold GAG types (**Supp. Fig. 3**). By day 21, there were more adherent monocyte-derived cells in response to MSC conditioned media generated in Heparin scaffolds versus either chondroitin sulfate scaffolds.

## 4. Discussion

The regeneration of complex CMF bone defects requires the interaction of biomaterial implants with multiple cell types. Previously, mineralized collagen scaffolds fabricated with CS6 glycosaminoglycan content have been shown to promote greater calcium and phosphorous mineral after 28 days of hMSC culture compared to CS4- and Heparin-containing scaffolds [65]. In order to successfully regenerate CMF defects, we must consider not only osteogenesis, but also mature vessel formation, resorption of the implant by osteoclasts, and the immune response. Here, our goal was to explore the potential role of scaffold GAG content, notably comparing chondroitin-6-sulfate, chondroitin-4-sulfate, and Heparin, on processes related to osteoclastogenesis, vasculogenesis, and immune response. By investigating the impact of glycosaminoglycans on other cells involved in bone repair, we hope to determine which GAG could have the most positive impact on CMF defect repair in biomaterials. To evaluate this goal, we compared the effect of the secretome generated by MSCs as a function of scaffold GAG content on subsequent activity of a model endothelial cell population (HUVECs) as a means for assessing angiogenic potential. Additionally, we evaluated patterns of monocyte specification using a THP-1 cell line capable of differentiating towards M1 and M2 macrophage phenotypes, which represent critical comparison regarding inflammatory response to injury, as well as towards the osteoclast lineage, which is responsible for bone remodeling.

Osteoclastogenesis is important for maintaining healthy bone homeostasis by controlling bone resorption. Although osteoclastogenesis is a necessary process in bone homeostasis, there is the potential to aid regenerative healing by transiently reducing osteoclast activity for a period of time immediately after biomaterial implantation. Osteoclast activity can be regulated by osteoprotegerin (OPG), which when produced by MSCs blocks RANKL-mediated RANK receptor activation on osteoclasts. Previously, work by *Ren et al*. has demonstrated that OPG produced by MSCs in mineralized collagen scaffolds reduced osteoclast activity [14, 15], and that MSCs transduced to boost OPG production can further limit bone resorption activity by osteoclasts [14]. However, for clinical translation it is advisable to avoid transduction of patient cells. Here, we examined the impact of scaffold glycosaminoglycans on osteoclasts and osteoclastogenesis as one of the cell types of interest in promoting healing of CMF defects. We found that hMSCs seeded on mineralized collagen-GAG scaffolds have significantly (p < 0.05) different OPG expression dependent on the GAG used, with Heparin contributing to the highest amount of OPG released at the later stages of the study (days 15-21). However, literature has noted that addition of Heparin to osteoblasts blocks the ability of OPG to bind to RANKL receptors [75]. This suggests that although Heparin scaffolds may enhance OPG production, it may not be functionally capable of reducing osteoclast activity. Future studies will directly target this relationship in order to define the role of scaffold GAG content on reciprocal signaling between osteoclasts and hMSCs in the context of hMSC biosynthetic and osteoclast degradative processes. While this study is confined to well-defined in vitro systems to investigate signaling crosstalk, future *in vivo* studies will be required to assess functional shifts in true regenerative potential in response to the presence of osrteoprogenitors, osteoclasts, and immune cells within the scaffold microenvironment.

We further examined osteoclast differentiation capacity of THP-1 monocytes in response to the MSC secretome generated on mineralized collagen scaffolds by assessing expression of two genes and two proteins associated with monocyte osteoclastogenesis. Expression of *SEMA4D*, an osteoclast-exclusive marker, was downregulated significantly (p < 0.05) in CS4 and Heparin at day 3 and in all groups on day 21. Similarly, *CTHRC1*, expressed by active osteoclasts, was downregulated significantly in CS6 and Heparin media at day 3. All scaffold groups seem to promote a MSC secretome that inhibits osteoclast gene expression, which is consistent with the osteoclast inhibitory role of increased OPG production. Notably, there were no differences in osteoclast protein expression of CT-1 and PDGF-BB in response to the secretome generated by MSCs regardless of scaffold GAG content. Both CT-1 and PDGF-BB were significantly (p < 0.05) downregulated in response to MSC secretome regardless of scaffold glycosaminoglycans compared to the basal media control, most significantly for the chondroitin groups. This downregulation suggests that the MSC secretome in response to the glycosaminoglycans used in mineralized collagen scaffolds did not promote osteoclast differentiation from monocytes. TRAP staining subsequently showed that after 14 and 21 days of culture, no visible sign of osteoclasts were found, though adherent cells were present in all samples, possibly indicating macrophage differentiation instead. Taken together, these data suggest scaffold glycosaminoglycans content may indirectly reduce osteoclast differentiation via factors produced endogenously by MSCs. The ability to shift the activity window of osteoclasts in and around a biomaterial implant during early versus later stages of healing could be advantageous and will require future efforts to compare functional metrics of osteoclast activity within the scaffolds. Notably, there are needs for quantitative assays to study co-cultures of MSCs and osteoclasts within a three-dimensional biomaterial to better determine scaffold design paradigms to alter osteoclast-mediated resorption and remodeling of tissue engineering scaffolds.

Mature vasculature formation is important for healthy bone and nutrient transport to continue new bone growth [76]. Here, we examined the role of MSC secreted factors as a function of scaffold GAG content on endothelial cell activity. Mesenchymal stem cells and osteoblasts are known to secrete VEGF to influence endothelial cell migration, proliferation, and differentiation [13, 77]. After 21 days of hMSC culture on mineralized collagen-GAG scaffolds, we discovered that VEGF secretion was significantly (p < 0.05) greater at days 9-15 in the CS6 scaffolds versus Heparin scaffolds. While earlier studies of hMSCs on mineralized collagen-GAG scaffolds have demonstrated protein release of angiogenin was significantly greater in the CS6 group compared to CS4 and Heparin groups [65], the effect of MSC secretome on endothelial cell activity had not previously been examined. When hMSC-GAG conditioned media was added to a Matrigel assay with HUVECs, there was significantly (p < 0.01) greater network length in CS6 scaffold conditioned media than all other groups. Though reduced, CS4 and Heparin conditioned media both promoted significantly (p < 0.05) greater network lengths than a non-conditioned media control. These results suggest the potential for scaffold GAG content to produce a hierarchy of factors able to promote angiogenic processes, most notably within scaffolds containing CS6. A possible explanation of CS6 having a greater impact on angiogenesis could be from its marine-derived nature, as marine-derived glycosaminoglycans have demonstrated better therapeutic potential than terrestrial glycosaminoglycans [78]. Interestingly, marine proteoglycans from sharks have shown to reduce matrix metalloproteinase expression to inhibit angiogenesis, but glycosaminoglycans from the same source did not have this effect [79]. While direct co-culture of endothelial cells with MSCs within scaffolds is possible [80], ongoing efforts will first pair hMSC-seeded scaffolds maintained in co-culture with hydrogels containing endothelial and stromal cells to study the effects of secreted factors on vessel maturation (e.g., network architecture; tight junction formation; basement membrane deposition) [81-84].

While macrophage plasticity is more complex than the traditional M1 vs. M2 phenotype comparison, the M1 to M2 transition is important for healing and regeneration of wounds, and without a proper transition persistent inflammation can cause fibrous tissue formation and healing limitations [68, 85, 86]. It is ideal for M1 macrophages to be present and active in the early stages of wound healing and eventually transition to M2 macrophages in the later stages of healing, and these timeframes usually range from days to weeks [87-89]. Literature has suggested that soluble Heparin has potential anti-inflammatory effects; however, the role of matrix immobilized heparan sulfate (HS) content may act via significantly different processes, leaving the opportunity to design *in vivo* studies to accurately determine this phenomena [90, 91]. Literature has also noted that chondroitin sulfate has anti-inflammatory effects by decreasing IL-1β and TNFα cytokine production in chondrocytes [92]. We investigated the ability of monocytes in hMSC-GAG conditioned media to differentiate towards M1 versus M2 macrophages over the course of 21 days. Notably, conditioned media generated by all scaffold groups had a lower expression of M1 associated gene *IL-1*β at days 3, 7, and 21 than the control.CS6 conditioned media had the greatest persistent upregulation of the M1-associated gene *TNF*α through days 7-14 with a dampening in expression at day 21. However, CS4 and Heparin induced a significant increase in *TNF*α expression at Day 21. This would suggest that CS6 conditioned media could allow for the macrophage temporal phenotypic transition from M1 to M2. CS6 conditioned media had the lowest protein expression compared to the control, significantly so with regards to pro-inflammatory proteins IFNγ, TNFα, and IL-1β. Overall, while hMSCs produced high amounts of IL-6 protein regardless of GAG type, scaffold glycosaminoglycan conditioned media drove limited pro-inflammatory phenotype in monocytes. In particular, CS6 scaffold conditioned media contributed to reduced IL6 production in monocytes, suggesting a possible anti-inflammatory effect.

In examining M2-associated markers, we observed that conditioned media generated by MSCs in scaffolds containing chondroitin sulfates drove upregulation of M2a gene *CCL22* at day 14, while CCL17 was upregulated in all groups at day 7 and 14.Additionally, gene expression of *CD163*, a M2c gene, was lower in all GAGs compared to the control at days 3, 7 and 21, and we observed a significant drop in expression of the M2c gene *MARCO* at day 7 in all groups as well. MSC mediated expression of pro-healing cytokines CCL18, CCL22, and IL-4 were significantly (p < 0.05) downregulated in all scaffold glycosaminoglycan groups. Together, these data suggest MSC conditioned media generated in mineralized collagen scaffolds, regardless of GAG content, largely downregulated M1 and M2 associated biomarkers, suggesting a potential immunosuppressive effect of the mineralized collagen scaffold. In the future, we plan to perform *in vivo* studies as well as seed M0 macrophages directly onto glycosaminoglycan-containing scaffolds. Overall differences in glycosaminoglycan properties could be due to sourcing differences (marine v. terrestrial), as marine-derived glycosaminoglycans have differences in sulfation leading to charge changes and can contain rare disaccharide units compared to terrestrial GAGs [78]. The marine CS6 that we implemented in this study has been extensively used and characterized in literature in the context of our scaffolds [40-44]. In the future, we plan to investigate the differences between marine and terrestrial CS6, CS4 and Heparin on these various cell types.

## 5. Conclusions

The goal of this study was to evaluate the impact of scaffold glycosaminoglycan content on the secretome generated by embedded MSCs, and the subsequent effect on osteoclastogenesis, angiogenesis, and immune processes essential for craniofacial bone regeneration. The direct effect of including GAGs in mineralized collagen scaffolds was previously investigated for osteogenic potential, with CS6 and Heparin scaffolds maximally promoting mineral formation *in vitro*. Here, we found that while inclusion of Heparin promoted the greatest release of OPG, all scaffolds downregulated osteoclast-associated protein expression. Further, scaffolds containing CS6 showed the greatest expression of VEGF as well as the most substantial endothelial tube formation, indicating increased angiogenic potential. Finally, conditioned media from all scaffold variants regardless of GAG content had an immunosuppresive effect that generated limited pro-and anti-inflammatory macrophage protein secretion, with CS6 scaffolds promoting the least amount of monocyte or macrophage IL6 production as well as the most substantial M1 to M2 gene expression transition. Taken together, these data suggest that scaffolds containing Heparin have the best potential to inhibit osteoclastogenesis, while those containing CS6 have the best potential to promote angiogenesis and mitigate the pro-inflammatory immune environment. Future directions include combining GAGs at different ratios and observing the presence of a synergistic versus antagonistic effect on the secretome of hMSCs. Furthermore, co-cultures of osteoclasts and macrophages, as well as *in vivo* experiments using pig models, which are one of the most relevant critical sized defect models, will be implemented to better understand the role of GAGs during bone regeneration.

## Acknowledgements

Research reported in this publication was supported by the National Institute of Dental and Craniofacial Research of the National Institutes of Health under Award Number R21 DE026582 (BACH) and R01 DE030491 (BACH). We are also grateful for the funding for this study provided by the NSF Graduate Research Fellowship (DGE-1144245 to MJD and MTN; DGE-1746047 to VK) and the Chemistry-Biology Interface Research Training Program at the University of Illinois (T32 GM070421, VK). The interpretations and conclusions presented are those of the authors and are not necessarily endorsed by the NIH or NSF.

The authors would like to acknowledge Dr. Bahaa Fadl-Alla and the College of Veterinary Medicine for assistance with real-time PCR. The authors would also like to acknowledge the Harley Lab for assistance with reviewing the manuscript and results. Additional support was provided by the Carl R. Woese Institute for Genomic Biology and the Chemical and Biomolecular Engineering Dept. at the University of Illinois at Urbana-Champaign.

## Data Availability

The raw and processed data required to reproduce these findings are available to download from Kolliopoulos, Vasiliki (2020) upon journal publication, “Data Repository: Glycosaminoglycan content of a mineralized collagen scaffold promotes mesenchymal stem cell secretion of factors to modulate angiogenesis and monocyte differentiation”, Mendeley Data, V1, doi: 10.17632/mpggxptxfb.1.

## Author Contributions

We describe contributions to the manuscript using the Contributor Roles Taxonomy (CRediT) [93, 94]: *Writing – Original Draft*: M.D. and V.K.; *Writing – Review & Editing:* M.D., V.K, M.N, and B.A.C.H.; *Conceptualization:* M.D., V.K, and B.A.C.H.; *Investigation:* M.D., V.K, and M.N; *Methodology:* M.D. and V.K; *Formal Analysis:* M.D. and V.K.; *Data Curation:* M.D. and V.K.; *Visualization:* M.D. and V.K.; *Project Administration:* B.A.C.H.; *Resources:* B.A.C.H.; *Funding Acquisition*: B.A.C.H.; Supervision: B.A.C.H.

## References

1. Lew, T.A., et al., Characterization of craniomaxillofacial battle injuries sustained by United States service members in the current conflicts of Iraq and Afghanistan. J Oral Maxillofac Surg, 2010. 68(1): p. 3–7.

2. Greenwald, A.S., Scott D. Boden, Victor M. Goldberg, Yusuf Khan, Cato T. Laurencin, and Randy N. Rosier, Bone-graft substitutes: facts, fictions, and applications. Journal of Bone and Joint Surgery, 2001. 83(2): p. 98–103.

3. Cg, F., Bone-grafting and bone-graft substitutes. Journal of Bone and Joint Surgery, 2002. 84(3): p. 454–464.

4. Kinaci, A., Valentin Neuhaus, and David C. Ring, Trends in bone graft use in the United States. Orthopedics, 2014. 37(9): p. e783–e788.

5. Xiao, D., et al., Estimating Reference Shape Model for Personalized Surgical Reconstruction of Craniomaxillofacial Defects. IEEE Trans Biomed Eng, 2021. 68(2): p. 362–373.

6. Costello, B.J., et al., Regenerative medicine for craniomaxillofacial surgery. Oral Maxillofac Surg Clin North Am, 2010. 22(1): p. 33–42.

7. Kinoshita, Y. and H. Maeda, Recent developments of functional scaffolds for craniomaxillofacial bone tissue engineering applications. ScientificWorldJournal, 2013. 2013: p. 863157.

8. Armiento, A.R., et al., Functional Biomaterials for Bone Regeneration: A Lesson in Complex Biology. Advanced Functional Materials, 2020: p. 1909874.

9. Pajarinen, J., et al., Mesenchymal stem cell-macrophage crosstalk and bone healing. Biomaterials, 2019. 196: p. 80–89.

10. Ignatius, A. and C. Sobacchi, Editorial: Innate Immunity in the Context of Osteoimmunology. Front Immunol, 2020. 11: p. 603.

11. Schlundt, C., et al., Immune modulation as a therapeutic strategy in bone regeneration. J Exp Orthop, 2015. 2(1): p. 1.

12. Lu, L.Y., et al., Pro-inflammatory M1 macrophages promote Osteogenesis by mesenchymal stem cells via the COX-2-prostaglandin E2 pathway. J Orthop Res, 2017. 35(11): p. 2378–2385.

13. Kyurkchiev, D., et al., Secretion of immunoregulatory cytokines by mesenchymal stem cells. World Journal of Stem Cells, 2014. 6: p. 552–570.

14. Ren, X., et al., Osteoprotegerin reduces osteoclast resorption activity without affecting osteogenesis on nanoparticulate mineralized collagen scaffolds. Science Advances, 2019. 5: p. 1–12.

15. Ren, X., et al., Nanoparticulate mineralized collagen glycosaminoglycan materials directly and indirectly inhibit osteoclastogenesis and osteoclast activation. Journal of tissue engineering and regenerative medicine, 2019. 13(5): p. 823–834.

16. Ono, T., et al., RANKL biology: bone metabolism, the immune system, and beyond. Inflamm Regen, 2020. 40: p. 2.

17. Takayanagi, H., Osteoimmunology - Bidirectional dialogue and inevitable union of the fields of bone and immunity. Proc Jpn Acad Ser B Phys Biol Sci, 2020. 96(4): p. 159–169.

18. Sohn, H.S. and J.K. Oh, Review of bone graft and bone substitutes with an emphasis on fracture surgeries. Biomater Res, 2019. 23: p. 9.

19. Wang, W. and K.W.K. Yeung, Bone grafts and biomaterials substitutes for bone defect repair: A review. Bioact Mater, 2017. 2(4): p. 224–247.

20. Faour, O., et al., The use of bone graft substitutes in large cancellous voids: any specific needs? Injury, 2011. 42 Suppl 2: p. S87–90.

21. Athanasiou VT P.D., Panagopoulos A, Saridis A, Scopa CD, Megas P., Histological comparison of autograft, allograft-DBM, xenograft, and synthetic grafts in a trabecular bone defect: An experimental study in rabbits. Med Sci Monit., 2010. 16(1): p. BR24–31.

22. Bauer, T.W., and George F. Muschler, Bone graft materials: an overview of the basic science. Clinical Orthopaedics and Related Research, 2000. 371: p. 10–27.

23. (CDC), C.f.D.C., Transmission of HIV through bone transplantation: case report and public health recommendations. MMWR. Morbidity and mortality weekly report, 1988. 37: p. 597.

24. Stevenson, S., and M. Horowitz, The response to bone allografts. JBJS, 1992. 74(6): p. 939–950.

25. Ren, X., et al., Nanoparticulate mineralized collagen scaffolds induce in vivo bone regeneration independent of progenitor cell loading or exogenous growth factor stimulation. Biomaterials, 2016. 89: p. 67–78.

26. Weisgerber, D.W., S.R. Caliari, and B.A. Harley, Mineralized collagen scaffolds induce hMSC osteogenesis and matrix remodeling. Biomater Sci, 2015. 3(3): p. 533–42.

27. Kanungo, B.P., et al., Characterization of mineralized collagen-glycosaminoglycan scaffolds for bone regeneration. Acta Biomater, 2008. 4(3): p. 490–503.

28. Cunniffe, G.M., et al., Development and characterisation of a collagen nano-hydroxyapatite composite scaffold for bone tissue engineering. Journal of Materials Science: Materials in Medicine, 2010. 21(8): p. 2293–2298.

29. Al-Munajjed, A.A., et al., Development of a biomimetic collagen-hydroxyapatite scaffold for bone tissue engineering using a SBF immersion technique. J Biomed Mater Res B Appl Biomater, 2009. 90(2): p. 584–91.

30. A. Al-Munajjed, J.G.a.F.O.B., Development of a collagen calcium-phosphate scaffold as a novel bone graft substitute. Stud Health Technol Inform., 2008. 133: p. 11–20.

31. Harley, B.A., et al., Design of a multiphase osteochondral scaffold. II. Fabrication of a mineralized collagen-glycosaminoglycan scaffold. J Biomed Mater Res A, 2010. 92(3): p. 1066–77.

32. Lee, J.C., et al., Optimizing collagen scaffolds for bone engineering: effects of cross-linking and mineral content on structural contraction and osteogenesis. The Journal of craniofacial surgery, 2015. 26(6): p. 1992.

33. Lyons, F.G., et al., Novel microhydroxyapatite particles in a collagen scaffold: a bioactive bone void filler? Clin Orthop Relat Res, 2014. 472(4): p. 1318–28.

34. Wang, X., et al., Restoration of a Critical Mandibular Bone Defect Using Human Alveolar Bone-Derived Stem Cells and Porous Nano-HA/Collagen/PLA Scaffold. Stem Cells Int, 2016. 2016: p. 8741641.

35. Mourao, P.A.S., Distribution of chondroitin 4–sulfate and chondroitin 6–sulfate in human articular and growth cartilage. Arthritis Rheuma, 1988. 31: p. 1028–1033.

36. O’Brien, F.J., et al., The effect of pore size on cell adhesion in collagen-GAG scaffolds. Biomaterials, 2005. 26(4): p. 433–41.

37. O’Brien, F.J., Brendan A. Harley, Ioannis V. Yannas, and Lorna Gibson, Influence of freezing rate on pore structure in freeze-dried collagen-GAG scaffolds. Biomaterials, 2004. 25(6): p. 1077–1086.

38. Harley, B.A., et al., Mechanical characterization of collagen-glycosaminoglycan scaffolds. Acta Biomater, 2007. 3(4): p. 463–74.

39. Harley, B.A., Hyung-Do Kim, Muhammad H. Zaman, Ioannis V. Yannas, Douglas A. Lauffenburger, and Lorna J. Gibson, Microarchitecture of three-dimensional scaffolds influences cell migration behavior via junction interactions. Biophysical journal 2008. 95(8): p. 4013–4024.

40. Caliari, S.R. and B.A. Harley, The effect of anisotropic collagen-GAG scaffolds and growth factor supplementation on tendon cell recruitment, alignment, and metabolic activity. Biomaterials, 2011. 32(23): p. 5330–40.

41. Hortensius, R.A. and B.A. Harley, The use of bioinspired alterations in the glycosaminoglycan content of collagen-GAG scaffolds to regulate cell activity. Biomaterials, 2013. 34(31): p. 7645–52.

42. Dewey, M.J., et al., Anisotropic mineralized collagen scaffolds accelerate osteogenic response in a glycosaminoglycan-dependent fashion. RSC Adv, 2020. 10(26): p. 15629–15641.

43. Caliari, S.R. and B.A. Harley, Collagen-GAG scaffold biophysical properties bias MSC lineage choice in the presence of mixed soluble signals. Tissue Eng Part A, 2014. 20(17-18): p. 2463–72.

44. Tiffany, A.S., et al., The inclusion of zinc into mineralized collagen scaffolds for craniofacial bone repair applications. Acta biomaterialia, 2019.

45. Bouvier, M., et al., Ultrastructural and immunocytochemical study of bone-derived cells cultured in three-dimensional matrices: influence of chondroitin-4 sulfate on mineralization. Differentiation, 1990. 45(2): p. 128–37.

46. Rajgopal, R., et al., The effects of heparin and low molecular weight heparins on bone. Thromb Res, 2008. 122(3): p. 293–8.

47. Ban, J.Y., S.W. Kang, and G.-J. Park, Heparin increases the osteogenic effect of recombinant human bone morphogenetic protein-2 in the rabbit bone defect model. Animal Cells and Systems, 2015. 19(5): p. 312–320.

48. Sun, B., et al., Crosslinking heparin to collagen scaffolds for the delivery of human platelet-derived growth factor. J Biomed Mater Res B Appl Biomater, 2009. 91(1): p. 366–72.

49. Chen, C., et al., Collagen/heparin sulfate scaffolds fabricated by a 3D bioprinter improved mechanical properties and neurological function after spinal cord injury in rats. J Biomed Mater Res A, 2017. 105(5): p. 1324–1332.

50. Tong M T.B., Hekking IM, Vermeij M, Barritault D, Van Neck JW, Stimulated neovascularization, inflammation resolution and collagen maturation in healing rat cutaneous wounds by a heparan sulfate glycosaminoglycan mimetic, OTR4120. Wound Repair Regen, 2009. 17(6): p. 840–852.

51. Kowitsch, A., G. Zhou, and T. Groth, Medical application of glycosaminoglycans: a review. J Tissue Eng Regen Med, 2018. 12(1): p. e23–e41.

52. Severin, I.C., et al., Glycosaminoglycan analogs as a novel anti-inflammatory strategy. Front Immunol, 2012. 3: p. 293.

53. Anderson, J. and S. Cramer, Perspectives on the Inflammatory, Healing, and Foreign Body Responses to Biomaterials and Medical Devices. 2015: p. 13–36.

54. BOGDAN-MIHAI NEAMŢU, a.A.B., Glycosaminoglycan-based biomaterials used in wound healing. AMT, 2019. 24(3): p. 84.

55. Lynn, A.K., et al., Design of a multiphase osteochondral scaffold. I. Control of chemical composition. J Biomed Mater Res A, 2010. 92(3): p. 1057–65.

56. Caliari, S.R., et al., The influence of collagen-glycosaminoglycan scaffold relative density and microstructural anisotropy on tenocyte bioactivity and transcriptomic stability. J Mech Behav Biomed Mater, 2012. 11: p. 27–40.

57. DeCicco-Skinner, K.L., et al., Endothelial cell tube formation assay for the in vitro study of angiogenesis. Journal of visualized experiments : JoVE, 2014(91): p. e51312–e51312.

58. Carpentier, G., et al., Angiogenesis Analyzer for ImageJ — A comparative morphometric analysis of “Endothelial Tube Formation Assay” and “Fibrin Bead Assay”. Scientific Reports, 2020. 10(1): p. 11568.

59. Maess, M.B., S. Sendelbach, and S. Lorkowski, Selection of reliable reference genes during THP-1 monocyte differentiation into macrophages. BMC Mol Biol, 2010. 11: p. 90.

60. University of Rochester Medical Center Rochester, N. Histology Forms and Protocols. TRAP staining for Paraffin Sections 2021; Available from: https://www.urmc.rochester.edu/musculoskeletal-research/core-services/histology/protocols.aspx.

61. Scott, R.A., K.L. Kiick, and R.E. Akins, Substrate stiffness directs the phenotype and polarization state of cord blood derived macrophages. Acta Biomater, 2020.

62. Taraballi, F., et al., Biomimetic collagenous scaffold to tune inflammation by targeting macrophages. J Tissue Eng, 2016. 7: p. 2041731415624667.

63. Dewey, M.J., et al., Shape-fitting collagen-PLA composite promotes osteogenic differentiation of porcine adipose stem cells. Journal of the Mechanical Behavior of Biomedical Materials, 2019. 95: p. 21–33.

64. Weisgerber, D.W., S.R. Caliari, and B.A.C. Harley, Mineralized collagen scaffolds induce hMSC osteogenesis and matrix remodeling. Biomaterials Science, 2015. 3: p. 533–542.

65. Dewey, M.J., et al., Anisotropic mineralized collagen scaffolds accelerate osteogenic response in a glycosaminoglycan-dependent fashion. RSC Advances, 2020. 10(26): p. 15629–15641.

66. Tiffany, A.S., et al., The inclusion of zinc into mineralized collagen scaffolds for craniofacial bone repair applications. Acta Biomaterialia, 2019. 93: p. 86–96.

67. Witherel, C.E., et al., Response of human macrophages to wound matrices in vitro. Wound Repair Regen, 2016. 24(3): p. 514–24.

68. Spiller, K.L., et al., Differential gene expression in human, murine, and cell line-derived macrophages upon polarization. Exp Cell Res, 2016. 347(1): p. 1–13.

69. Huleihel, L., Dziki, J.L., Bartolacci, J.G., Rausch, T., Scarritt, M.E., Cramer, M.C., Vorobyov, T., LoPresti, S.T., Swineheart, I.T., White, L.J. and Brown, B.N., Macrophage phenotype in response to ECM bioscaffolds. Seminars in immunology, 2017. 29: p. 2–13.

70. Tugal, D., Xudong Liao, and Mukesh K. Jain, Transcriptional control of macrophage polarization. Arteriosclerosis, thrombosis, and vascular biology, 2013. 33(6): p. 1135–1144.

71. Spiller, K.L., D.O. Freytes, and G. Vunjak-Novakovic, Macrophages modulate engineered human tissues for enhanced vascularization and healing. Ann Biomed Eng, 2015. 43(3): p. 616–27.

72. Genin, M., et al., M1 and M2 macrophages derived from THP-1 cells differentially modulate the response of cancer cells to etoposide. BMC Cancer, 2015. 15: p. 577.

73. Mantovani, A., et al., The chemokine system in diverse forms of macrophage activation and polarization. Trends Immunol, 2004. 25(12): p. 677–86.

74. Han, Y., et al., Paracrine and endocrine actions of bone-the functions of secretory proteins from osteoblasts, osteocytes, and osteoclasts. Bone Res, 2018. 6: p. 16.

75. Irie, A., et al., Heparin enhances osteoclastic bone resorption by inhibiting osteoprotegerin activity. Bone, 2007. 41(2): p. 165–74.

76. Ngo, M.T. and B.A.C. Harley, Angiogenic biomaterials to promote therapeutic regeneration and investigate disease progression. Biomaterials, 2020. 255: p. 120207.

77. Hu, K. and B.R. Olsen, The roles of vascular endothelial growth factor in bone repair and regeneration. Bone, 2016. 91: p. 30–38.

78. Valcarcel, J., et al., Glycosaminoglycans from marine sources as therapeutic agents. Biotechnol Adv, 2017. 35(6): p. 711–725.

79. Kobayashi, T., et al., Chondroitin sulfate proteoglycans from salmon nasal cartilage inhibit angiogenesis. Biochem Biophys Rep, 2017. 9: p. 72–78.

80. Duffy, G.P., et al., Bone marrow-derived mesenchymal stem cells promote angiogenic processes in a time-and dose-dependent manner in vitro. Tissue Eng Part A, 2009. 15(9): p. 2459–70.

81. Ngo, M.T. and B.A.C. Harley, Perivascular signals alter global gene expression profile of glioblastoma and response to temozolomide in a gelatin hydrogel. Biomaterials, 2019. 198: p. 122–134.

82. Ngo, M.T. and B.A.C. Harley, The influence of hyaluronic acid and glioblastoma cell co-culture on the formation of endothelial cell networks in gelatin hydrogels. Adv Healthc Mater, 2017. 6(21): p. 1700687.

83. Ngo, M.T., et al., Hydrogels containing gradients in vascular density reveal dose-dependent role of angiocrine cues on stem cell behavior. in revision, 2021.

84. Barnhouse, V., et al., Perivascular secretome influences hematopoietic stem cell maintenance in a gelatin hydrogel. Ann Biomed Eng, 2021. 49(2): p. 780–792.

85. Witherel, C.E., et al., Regulation of extracellular matrix assembly and structure by hybrid M1/M2 macrophages. Biomaterials, 2021. 269: p. 120667.

86. Witherel, C.E., et al., Immunomodulatory effects of human cryopreserved viable amniotic membrane in a pro-inflammatory environment in vitro. Cell Mol Bioeng, 2017. 10(5): p. 451–462.

87. Spiller, K.L., et al., Sequential delivery of immunomodulatory cytokines to facilitate the M1-to-M2 transition of macrophages and enhance vascularization of bone scaffolds. Biomaterials, 2015. 37: p. 194–207.

88. Weingarten, M.S., et al., Response of human macrophages to wound matrices in vitro. Wound Repair and Regeneration, 2016. 24: p. 514–524.

89. Spiller, K.L., D.O. Freytes, and G. Vunjak-Novakovic, Macrophages Modulate Engineered Human Tissues for Enhanced Vascularization and Healing. Annals of Biomedical Engineering, 2015. 43: p. 616–627.

90. Jenmalm, M.C., Effects of low molecular weight heparin on the polarization and cytokine profile of macrophages and T helper cells in vitro. Nature Scientific Reports, 2018. 4166: p. 2–10.

91. Mousavi, S., et al., Anti-Inflammatory Effects of Heparin and Its Derivatives : A Systematic Review. Advances in Pharmacological Sciences, 2015. 2015: p. 1–14.

92. García, A.G., J. Vergés, and E. Montell, Immunomodulatory and anti-inflammatory effects of chondroitin sulphate. J Cell Mol Med, 2009. 13: p. 1451–1463.

93. Brand, A., et al., Beyond authorship: attribution, contribution, collaboration, and credit. Learned Publishing, 2015. 28(2): p. 151–155.

94. Allen, L., et al., Publishing: Credit where credit is due. Nature, 2014. 508(7496): p. 312–3.

