## Supplementary material for "Glycosaminoglycan content of a mineralized collagen scaffold promotes mesenchymal stem cell secretion of factors to modulate angiogenesis and monocyte differentiation": Figure captions

**Fig. 1.** Experimental Outline. Mineralized collagen scaffolds were fabricated with glycosaminoglycans chondroitin-6-sulfate, chondroitin-4-sulfate, or Heparin. Human mesenchymal stem cells were seeded on scaffolds and conditioned media surrounding the scaffold was collected after 6 and 21 days. Conditioned media from scaffolds after 6 days were then used in a Matrigel tube formation assay with human umbilical cord endothelial cells to quantify tube formation after 6 and 12hrs of cell culture. Conditioned media from scaffolds after 21 days was assessed for OPG and VEGF release to determine possible effects on osteoclastogenesis and angiogenesis. Conditioned media were also used on human THP-1 monocytes for up to 21 days to assess differentiation into osteoclasts and M1 or M2 macrophages.

**Fig. 2.** Osteoprotegerin (OPG) and vascular endothelial growth factor (VEGF) cumulative release from hMSC-seeded mineralized collagen scaffolds containing glycosaminoglycans chondroitin-6-sulfate (CS6), chondroitin-4-sulfate (CS4), or heparin (Heparin). (A) Mesenchymal stem cells produce OPG to block RANKL on osteoblasts. RANK binding of osteoclast progenitors to RANKL signals osteoclast progenitors to differentiate into osteoclasts and promotes osteoclastogenesis. * denotes that the Heparin group has significantly (p < 0.05) greater release of OPG than both CS6 and CS4. (B) Mesenchymal stem cells and osteoblasts produce VEGF that acts on endothelial cells to promote angiogenesis by endothelial cell migration, proliferation, and differentiation. ^ indicates the CS6 group has significantly (p < 0.05) greater release of VEGF than the Heparin group. Data represented as average ± standard deviation (n=6).

**Fig. 3.** A Matrigel tube formation assay with HUVECs (6hr) was used to determine the influence of hMSC-glycosaminoglycan (chondroitin-6-sulfate (CS6), chondroitin-4-sulfate (CS4), and heparin (Heparin)) conditioned media on angiogenesis. Representative images of tube formation are presented on the left (Scale bar represents 0.2 mm and cells are false colored red for contrast), and analysis of images is presented on the right. Total network length was quantified with ImageJ and asterisks represent *p<0.05, **p<0.01, ***p<0.001 compared to Negative Control Media (basal hMSC media). Carrots (^^) represent p<0.01 compared to CS4, and hashtags (##) represent p<0.01 compared to Heparin. Data represented as mean (center line) with standard deviation (whiskers) (n=6).

**Fig. 4.** Macrophage phenotype-related gene expression of monocytes conditioned in MSC-conditioned media in response to scaffold glycosaminoglycan content (chondroitin-6-sulfate, CS6; chondroitin-4-sulfate, CS4; heparin, Heparin). Control group represents basal hMSC media with no hMSC conditioned factors combined with RPMI media on monocytes. Gene expression is represented as fold change compared to monocyte gene expression before media conditioning. M1 (pro-inflammatory) genes include IL1β and TNFα, M2a (anti-inflammatory) genes include CCL22 and CCL17, and M2c (anti-inflammatory) genes include CD163 and MARCO. Different letters on the same day correspond to significant (p < 0.05) differences between groups. Data represented as average ± standard deviation (n=5).

**Fig. 5**. Immune and angiogenesis-related cytokine secretion by mesenchymal stem cells cultured in scaffolds as a function of glycosaminoglycan (chondroitin-6-sulfate, CS6; chondroitin-4-sulfate, CS4; heparin, Heparin) content as well as monocytes subsequently cultured in MSC-conditioned media. Protein release is represented as a fold change compared to blank media control, while the control group represents monocytes in unconditioned media. (A) IL6 and VEGF fold change normalized to blank media control. Solid color represents hMSC contribution to cytokine release and patterned color represents monocyte contribution to cytokine release. White circles represent hMSC data points and black circles represent monocyte data points. * indicates the hMSC and monocyte contributions to IL6 were significantly (p < 0.05) different for the CS6 group. Data expressed as average (n=3). (B) Release of cytokines TNFα (M1) and CCL22 (M2) from monocytes cultured in GAG conditioned media compared to a control group cultured in normal media. * indicates secretion in the control group is significantly (p < 0.05) greater than the indicated group(s). Data expressed as average ± standard deviation (n=3).

**Fig. 6.** Osteoclast-related cytokine release and gene expression of monocytes conditioned in media from various glycosaminoglycans (chondroitin-6-sulfate, CS6; chondroitin-4-sulfate, CS4; heparin, Heparin). (A) Gene expression is represented as a fold change compared to monocyte gene expression before media conditioning. Osteoclast genes include *CTHRC1* and *SEMA4D*. Different letters on the same day correspond to significant (p < 0.05) differences between groups. Data represented as average ± standard deviation (n=5). (B) Protein release (CT-1 and PDGF-BB) is represented as a fold change compared to blank media control, with the control group representing monocytes in unconditioned media. * indicates control is significantly (p < 0.05) greater than other indicated groups. Data expressed as average ± standard deviation (n=3).
