## Supplemental material for "Glycosaminoglycan content of a mineralized collagen scaffold promotes mesenchymal stem cell secretion of factors to modulate angiogenesis and monocyte differentiation"

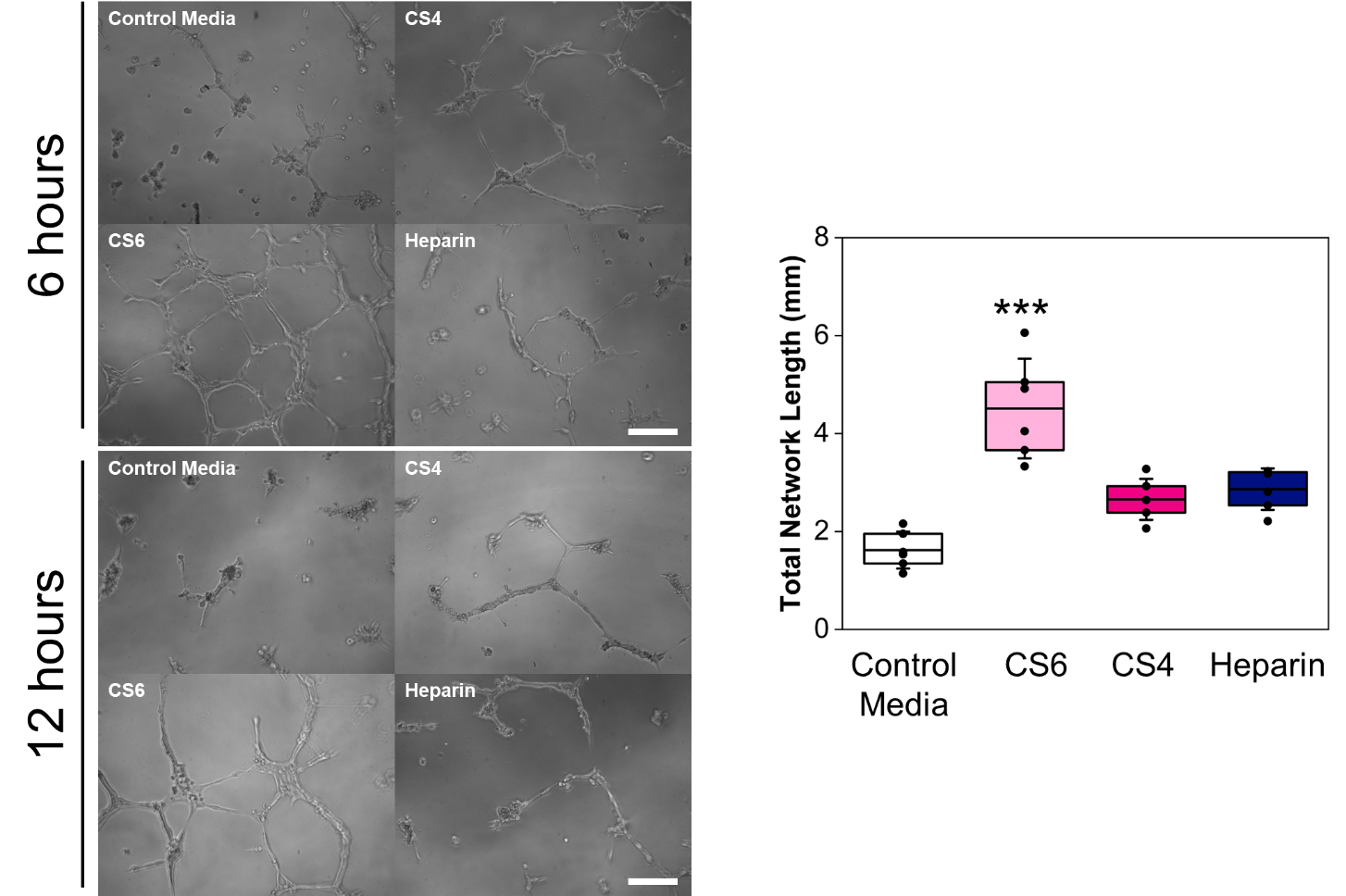

**Supp. Fig. 1.** A Matrigel tube formation assay with HUVECs (6 and 12hr) was used to determine the influence of hMSC-glycosaminoglycan (chondroitin-6-sulfate, CS6; chondroitin-4-sulfate, CS4; heparin, Heparin) conditioned media on angiogenesis. Negative control media group represents basal hMSC media. Representative images of tube formation are presented on the left (scale bar represents 0.2 mm), and analysis of 12 hr images is presented on the right. Total network length was quantified with ImageJ and asterisks represent ***p<0.001 compared to Negative Control. Data represented as mean (center line) with standard deviation (whiskers) (n=6).

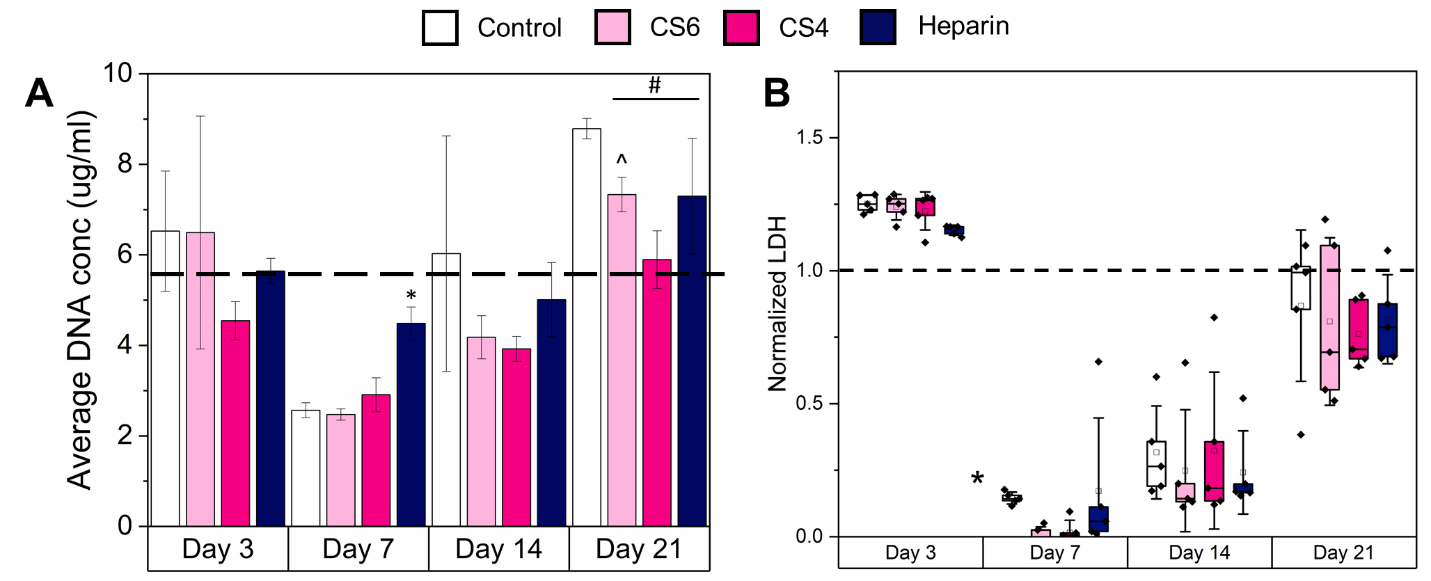

**Supp. Fig. 2** Cell viability of monocytes cultured in hMSC-GAG media (chondroitin-6-sulfate, CS6; chondroitin-4-sulfate, CS4; heparin, Heparin). Monocytes were cultured in control media (no GAG), CS6 media (CS6 hMSC-GAG), CS4 media (CS4 hMSC-GAG), and Heparin media (Heparin hMSC-GAG) for 21 days. Cell viability was determined by a Quant-iT™ PicoGreen® assay to observe average DNA concentrations, and a lactate dehydrogenase (LDH) assay was used to determine cytotoxicity. (A) An average DNA concentration of 5.79 µg/mL represents the activity of 1 million cells seeded into every well and is represented by the dashed line. * indicates one group was significantly (p < 0.05) different from the other groups on the same day, # indicates one group was significantly (p < 0.05) different from the control group of that day, ^ indicates one group was significantly (p < 0.05) different from CS4 of that day. Data represented as average ± standard deviation (n=5). (B) A LDH assay was used to determine the cytotoxicity of hMSC-GAG media on monocytes. LDH is released upon damage to cellular membranes and was recorded as luminescence and normalized to cells prior to adding to wells with conditioned media (a value of 1). Higher values indicate more LDH released and potentially greater cytotoxicity. * indicates one group was significantly (p < 0.05) different from the other groups on the same day. Data represented as average ± standard deviation (n=5).

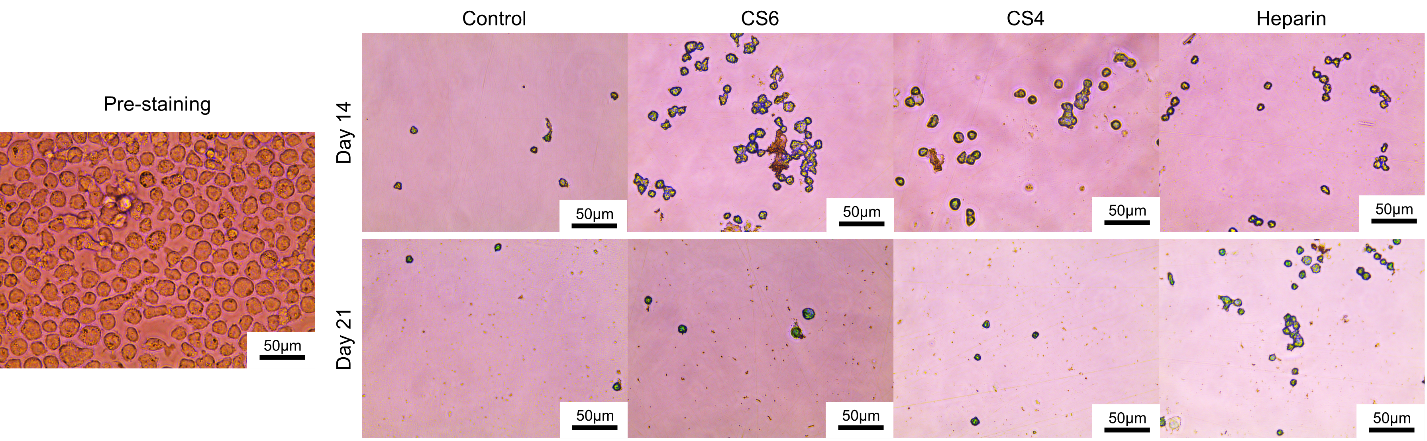

**Supp Fig. 3.** Tartrate-resistant acid phosphatase (TRAP) staining of THP-1 monocytes in hMSC-GAG conditioned media (chondroitin-6-sulfate, CS6; chondroitin-4-sulfate, CS4; heparin, Heparin). Images of cells before staining were taken and there was no distinguishable difference between groups, and cells were uniformly distributed throughout wells. After removing media, washing, and staining with TRAP, the remaining cells were imaged. Osteoclasts are stained in red, and other cell types are stained in green. Scale bar is 50 µm (n=6).

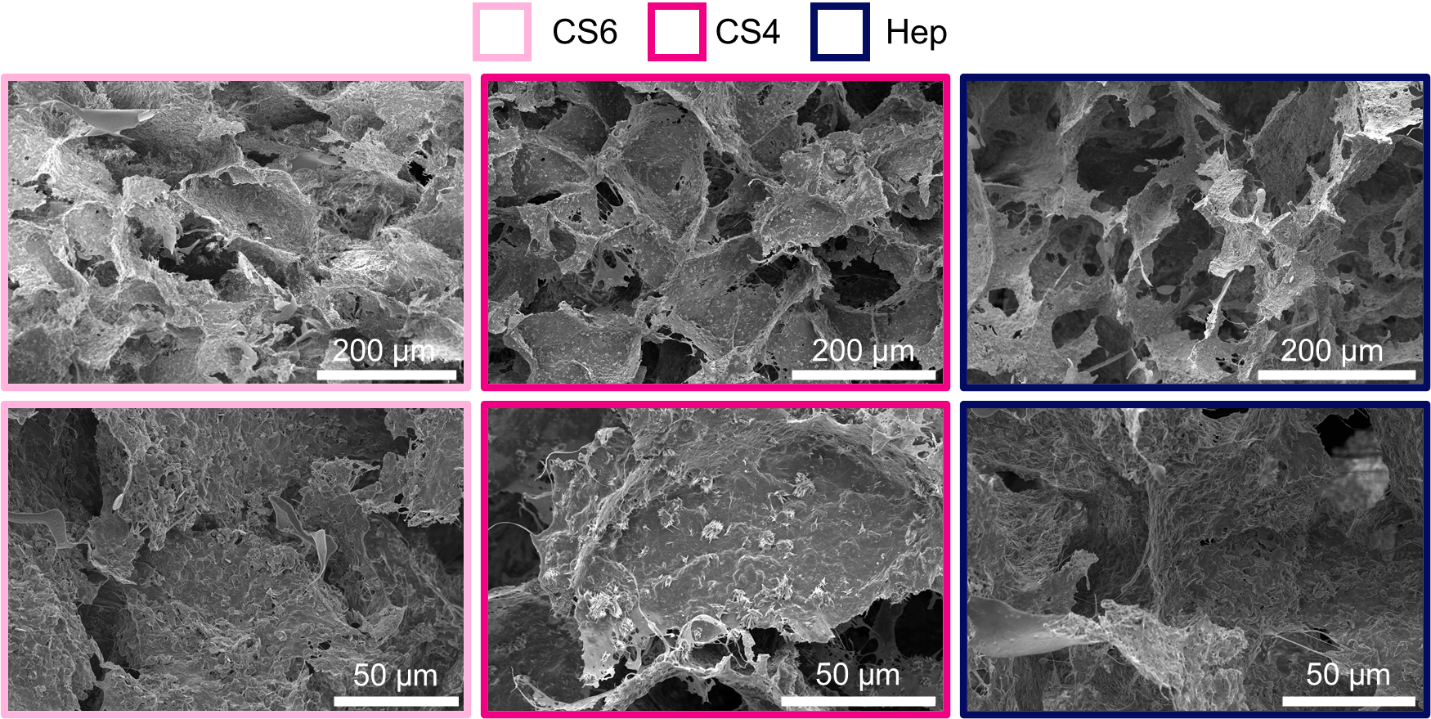

**Supp Fig. 4.** Scanning electron microscopy (SEM) representative images of the mineralized collagen-GAG scaffolds containing chondroitin-6-sulfate, CS6; chondroitin-4-sulfate, CS4; or heparin, Heparin.

| **Function** | **Gene symbol** | **Manufacturer** | **Assay ID** |
| --- | --- | --- | --- |
| Housekeeping | *h-ACTB* | TaqMan from LIFE TECHNOLOGIES CORPORATION | Hs01060665_g1 |
| M1 | *h-IL1B* |  | Hs01555410_m1 |
| M1 | *h-TNF* |  | Hs00174128_m1 |
| M2 | *h-MARCO* |  | Hs00198935_m1 |
| M2 | *h-CCL22* |  | Hs01574247_m1 |
| M2 | *h-CD163* |  | Hs00174705_m1 |
| M2 | *h-CCL17* |  | Hs00171074_m1 |
| Osteoclast | *h-CTHRC1* |  | Hs00298917_m1 |
| Osteoclast | *h-SEMA4D* |  | Hs00925667_m1 |

**Supp. Table 1** TaqMan genes used for RT-PCR to analyze M1 (pro-inflammatory), M2 (pro-healing) and osteoclast gene expression in THP-1 monocytes.

|  | | **Mesenchymal Stem Cell Cytokine Expression** | | |
| --- | --- | --- | --- | --- |
|  |  | CS4 | CS6 | Hep |
| M1 | IFN-gamma | 1.1 ± 0.38 | 0.76 ± 0.20 | 0.81 ± 0.19 |
|  | TNFa | 1.1 ± 0.23 | 0.78 ± 0.21 | 0.85 ± 0.17 |
|  | IL-1B | 1.1 ± 0.84 | 0.79 ± 0.31 | 0.79 ± 0.53 |
|  | IL-6 | 33 ± 15 | 31 ± 6.4 | 29 ± 6.8 |
| M2 | IL-10 | 1.2 ± 0.27 | 1.0 ± 0.20 | 0.98 ± 0.18 |
|  | CCL18 | 1.1 ± 0.32 | 0.78 ± 0.17 | 0.84 ± 0.17 |
|  | CCL22 | 1.1 ± 0.34 | 0.76 ± 0.21 | 0.83 ± 0.18 |
|  | MMP9 | 1.1 ± 0.30 | 0.78 ± 0.21 | 0.82 ± 0.16 |
|  | CCL17 | 1.1 ± 0.21 | 0.79 ± 0.23 | 0.92 ± 0.27 |
|  | IL-4 | 1.1 ± 0.40 | 0.72 ± 0.27 | 0.82 ± 0.28 |
|  | IL-13 | 1.06 ± 0.21 | 0.84 ± 0.21 | 0.88 ± 0.17 |
| Osteoclast | CT-1 | 1.2 ± 0.34 | 0.86 ± 0.20 | 0.86 ± 0.15 |
|  | PDGF-BB | 1.1 ± 0.29 | 0.82 ± 0.18 | 0.85 ± 0.15 |
| Vasculature | VEGF | 2.6 ± 0.52 | 1.7 ± 0.21 | 1.7 ± 0.11 |

**Supp. Table 2** Cytokine release profiles of hMSCs seeded on GAG-containing mineralized collagen scaffolds (chondroitin-6-sulfate, CS6; chondroitin-4-sulfate, CS4; heparin, Heparin) measured via a custom cytokine array (RayBiotech, Georgia, USA). Expression is represented as a fold change compared to blank media control, and no significant differences were found between groups. Data is colored as red representing a fold change of 0, yellow as a fold change of 1, and green as a fold change of 5. Data expressed as average ± standard deviation (n=3).

|  | | **Monocyte Cytokine Expression** | | | |
| --- | --- | --- | --- | --- | --- |
|  |  | Control | CS4 | CS6 | Hep |
| M1 | IFN-gamma | 1.1 ± 0.43 | 0.09 ± 0.43* | 0.11 ± 0.11* | 0.17 ± 0.29 |
|  | TNFa | 1.0 ± 0.40 | 0.46 ± 0.37 | 0.06 ± 0.11* | 0.20 ± 0.30 |
|  | IL-1B | 1.0 ± 1.4 | 0.41 ± 3.5 | 0.06 ± 1.6* | 0.31 ± 1.5 |
|  | IL-6 | 6.0 ± 5.2 | 0.95 ± 1.6 | 0.0 ± 0.0 | 2.3 ± 4.1 |
| M2 | IL-10 | 1.1 ± 0.49 | 0.46 ± 0.41 | 0.18 ± 0.31 | 0.48 ± 0.23 |
|  | CCL18 | 1.0 ± 0.41 | 0.24 ± 0.21* | 0.13 ± 0.17* | 0.22 ± 0.19* |
|  | CCL22 | 1.5 ± 0.61 | 0.41 ± 0.37* | 0.25 ± 0.25* | 0.47 ± 0.29* |
|  | MMP9 | 1.3 ± 0.69 | 0.28 ± 0.26 | 0.23 ± 0.21 | 0.43 ± 0.29 |
|  | CCL17 | 0.95 ± 0.39 | 0.52 ± 0.62 | 0.039 ± 0.068 | 0.22 ± 0.34 |
|  | IL-4 | 0.22 ± 0.089 | 0.0 ± 0.0* | 0.0 ± 0.0* | 0.0 ± 0.0* |
|  | IL-13 | 1.0 ± 0.41 | 0.43 ± 0.40 | 0.10 ± 0.17* | 0.28 ± 0.25 |
| Osteoclast | CT-1 | 1.2 ± 0.49 | 0.27 ± 0.23* | 0.21 ± 0.32* | 0.31 ± 0.26 |
|  | PDGF-BB | 1.1 ± 0.42 | 0.22 ± 0.21* | 0.15 ± 0.21* | 0.19 ± 0.15* |
| Vasculature | VEGF | 3.9 ± 0.64 | 3.8 ± 0.15 | 2.1 ± 0.19 | 3.8 ± 0.17 |

**Supp. Table 3** Cytokine release profiles of monocytes in hMSC-GAG conditioned media (chondroitin-6-sulfate, CS6; chondroitin-4-sulfate, CS4; heparin, Heparin) measured via a custom cytokine array (RayBiotech, Georgia, USA). Expression is represented as a fold change compared to blank media control, while “control” group represents monocytes in unconditioned media. * indicates significant (p < 0.05) difference in GAG group indicated compared to control for same cytokine. Data is colored as red representing a fold change of 0, yellow as a fold change of 1, and green as a fold change of 5. Data expressed as average ± standard deviation (n=3).
